# Effects of common mutations in the SARS-CoV-2 Spike RBD domain and its ligand the human ACE2 receptor on binding affinity and kinetics

**DOI:** 10.1101/2021.05.18.444646

**Authors:** Michael I. Barton, Stuart MacGowan, Mikhail Kutuzov, Omer Dushek, Geoffrey J. Barton, P. Anton van der Merwe

## Abstract

The interaction between the SARS-CoV-2 virus Spike protein receptor binding domain (RBD) and the ACE2 cell surface protein is required for viral infection of cells. Mutations in the RBD domain are present in SARS-CoV-2 variants of concern that have emerged independently worldwide. For example, the more transmissible B.1.1.7 lineage has a mutation (N501Y) in its Spike RBD domain that enhances binding to ACE2. There are also ACE2 alleles in humans with mutations in the RBD binding site. Here we perform a detailed affinity and kinetics analysis of the effect of five common RBD mutations (K417N, K417T, N501Y, E484K and S477N) and two common ACE2 mutations (S19P and K26R) on the RBD/ACE2 interaction. We analysed the effects of individual RBD mutations, and combinations found in new SARS-CoV-2 variants first identified in the UK (B.1.1.7), South Africa (B.1.351) and Brazil (P1). Most of these mutations increased the affinity of the RBD/ACE2 interaction. The exceptions were mutations K417N/T, which decreased the affinity. Taken together with other studies, our results suggest that the N501Y and S477N mutations primarily enhance transmission, the K417N/T mutations facilitate immune escape, and the E484K mutation facilitates both transmission and immune escape.

## Introduction

Since its identification in 2019, the second coronavirus able to induce a severe acute respiratory syndrome in humans, SARS-CoV-2, has resulted in the most severe global pandemic in 100 years. To date more than 135 million people have been infected, resulting in the deaths from the resulting disease, COVID-19, of more than 3 million people (“WHO Coronavirus (COVID-19) Dashboard,” 2021), and measures introduced to control spread have had harmful social and economic impacts. Fortunately, effective vaccines have been developed, and a global vaccination programme is underway (Mahase, 2021). New SARS-CoV-2 variants of concern are emerging that are making containment of the pandemic more difficult, by increasing transmissivity of the virus (Davies and Edmunds, 2021; Korber et al., 2020; Volz et al., 2021a, 2021b; Washington et al., 2021) and/or its resistance to protective immunity induced by previous infection or vaccines (Darby and Hiscox, 2021; Dejnirattisai et al., 2021; Garcia-Beltran et al., 2021; Madhi et al., 2021a, 2021b; Mahase, 2021).(Volz et al., 2021a, 2021b)

The SARS-CoV-2 virus enters cells following an interaction between the Spike (S) protein on its surface with angiotensin-converting enzyme 2 (ACE2) on cell surfaces (V’kovski et al., 2021). The receptor binding domain (RBD) of the Spike protein binds the membrane-distal portion of the ACE2 protein. The S protein forms a homotrimer, which is cleaved shortly after synthesis into two fragments that remain associated non-covalently: S1, which contains the RBD, and S2, which mediates membrane fusion following the binding of Spike to ACE2 (V’kovski et al., 2021). During the pandemic mutations have appeared in the Spike protein that apparently increase transmissivity (Davies and Edmunds, 2021; Korber et al., 2020; Volz et al., 2021a, 2021b; Washington et al., 2021). One that emerged early in Europe, D614G, and quickly became dominant globally (Korber et al., 2020), increases the density of intact Spike trimer on the virus surface by preventing premature dissociation of S1 from S2 following cleavage (Zhang et al., 2021, 2020). A later mutant, N501Y, which has appeared in multiple lineages, lies within the RBD domain, and increases its affinity for ACE2 (Starr et al., 2020; Supasa et al., 2021). These findings suggest that mutations that directly or indirectly enhance Spike binding to ACE2 will increase transmissivity.

Prior infection by SARS-CoV-2 and current vaccines induce antibody responses to the Spike protein, and most neutralizing antibodies appear to bind to the Spike RBD domain (Garcia-Beltran et al., 2021; Greaney et al., 2021a; Rogers et al., 2020). Some variants of concern have mutations in their RBD domain that confer resistance to neutralizing antibodies (Darby and Hiscox, 2021; Dejnirattisai et al., 2021; Garcia-Beltran et al., 2021; Madhi et al., 2021a, 2021b; Mahase, 2021). What is less clear is the precise effect of these mutations on the affinity and kinetics of the binding of RBD to ACE2. Previous studies of the interaction between the Spike RBD and ACE2 have produced a wide range of affinity and kinetic estimates under conditions (e.g. temperature) that are not always well defined (Lei et al., 2020; Shang et al., 2020; Supasa et al., 2021; Wrapp et al., 2020; Zhang et al., 2021, 2020). Precise information is needed to assess the extent to which RBD mutations have been selected because they enhance ACE2 binding or facilitate immune evasion.

In this study we undertook a detailed affinity and kinetic analysis of the interaction between Spike RBD and ACE2 at physiological temperatures, taking care to avoid common pitfalls. We used this optimized approach to analyse the effect of important common mutations identified in variants of RBD and ACE2. Both mutations of ACE2 (S19P, K26R) and most of the mutations of RBD (N501Y, E484K, and S477N) enhanced the interaction, with some RBD mutations (N501Y) increasing the affinity by ~10 fold. Increased binding was the result of decreases in dissociation rate constants (N501Y, S477N) and/or increases in association rate constants (N501Y, E484K). Although the K417N/T mutations found in the South African (B.1.351) and Brazilian (P.1) variants both decreased the affinity, the affinity-enhancing N501Y and E484K mutations that are also present in both variants confer a net ~4 fold increase in the affinity of their RBD domains for ACE2.

## Results

### Selection of variants

The focus of this study was to analyse common and therefore important variants of RBD and ACE2. Henceforth we will refer to the common ACE2 allele and RBD of the original SARS-CoV-2 strain sequenced in Wuhan as wild-type (WT). We chose mutations of RBD within the ACE2 binding site that have appeared independently in multiple SARS-CoV-2 lineages/clades (Fig. 1 and Fig. S1) (Hodcroft, 2021; Rambaut et al., 2020), suggesting that they confer a selective advantage, rather than emerged by chance, such as through a founder effect. The N501Y mutation has appeared in the B.1.1.7 (20I/501Y.V1), B.1.351 (20H/501Y.V2), and P.1 lineages (20J/501Y.V3) first identified in the UK, South Africa and Brazil, respectively. The E484K mutation is present in the B.1.351 and P.1 lineages and has appeared independently in many other lineages, including P.2 (20B/S.484K), B.1.1.318, B.1.525 (20A/S:4.4K), and B.1.526 (20C/S.484K). E484K has also appeared in VOC-202102-02, a subset of the B.1.1.7 lineage identified in the UK (“SARS-CoV-2 Variants of concern and variants under investigation - GOV.UK,” 2021). The S477N mutation became dominant for periods in Australia (clade 20F) and parts of Europe (20A.EU2), and then appeared in New York in lineage B.1.526 (H. Zhou et al., 2021). Mutations of K417 have appeared independently in the South African B.1.351 and Brazilian P.1 lineages. Interestingly, N501Y, E484K and S477N were the main mutations that appeared following random RBD mutagenesis and in vitro selection of mutants with enhanced ACE2 binding (Zahradník et al., 2021).

**Figure 1.**
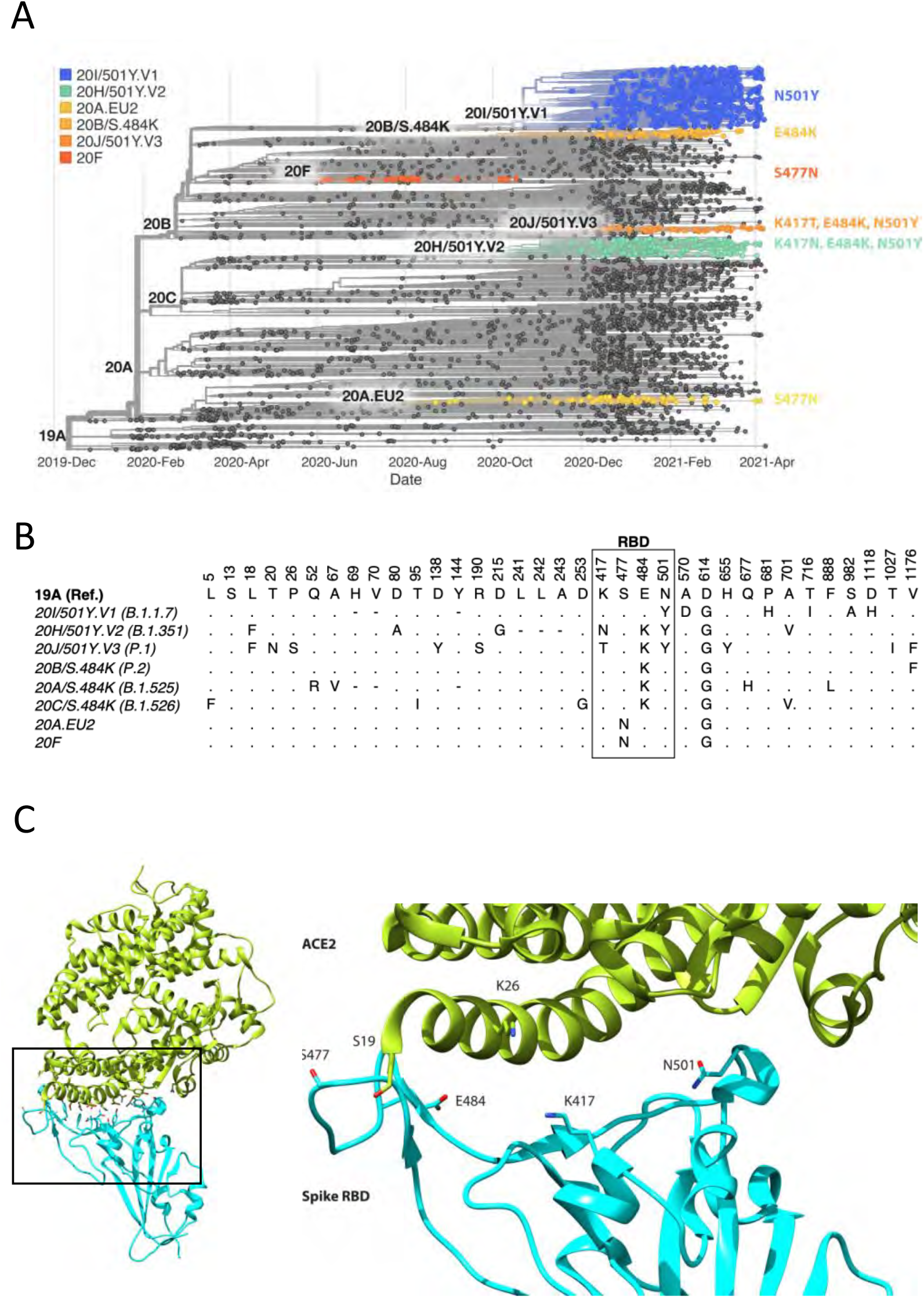
Spike RBD and ACE2 variants analysed in this study. (A) Phylogenetic tree illustrating the clades containing the RBD mutations investigated in this study. Constructed using TreeTime (Sagulenko et al., 2018) from the Nextstrain Global (Hadfield et al., 2018) sample of SARS-CoV-2 sequences from the GISAID database (Shu and McCauley, 2017) (Accessed 15th April 2021, N = 4,017). (B) Alignment illustrating the Spike residues that differ between SARS-CoV-2 variants, with the RBD mutants boxed. The variants are labelled with their clade designation from Nextstrain (Hadfield et al., 2018) and/or PANGO lineage (Rambaut et al., 2020) where relevant. The RBD mutations were collated from CoVariants (Hodcroft, 2021) and Nextstrain. (C) The structure of human ACE2 (green) in complex with SARS-CoV-2 Spike RBD (cyan). The area enclosed by the box is shown enlarged on the right, with the residues mutated in this study labelled. Drawn using UCSF Chimera (Pettersen et al., 2004) using coordinates from PDB 6m0j (Lan et al., 2020).

We selected for analysis the two most common mutations of ACE2 within the RBD binding site, K26R and S19P (Fig. 1C). They are present in 0.4% and 0.03%, respectively, of all samples in the gnomAD database (Karczewski et al., 2020), while other ACE2 mutations in the RBD binding site are much less frequent (<0.004%) (MacGowan et al., 2021). K26R is observed in all the major gnomAD populations but is most common in Ashkenazi Jews (1%), and (non-Finnish) north-western Europeans (0.6%). It is less common in Africans/African-Americans and South Asians (0.1%) and rare in Finnish (0.05%) and East-Asian (0.001%) populations. The S19P mutant is almost exclusively found in Africans/African-Americans (0.3 %).

### Measurement of affinity and kinetics

To measure the effects of these mutations on the affinity and kinetics of the RBD/ACE2 interaction we used surface plasmon resonance, which allows very accurate measurements, provided that common pitfalls are avoided, particularly protein aggregation, mass-transport limitations and rebinding (van der Merwe and Barclay, 1996; Myszka, 1997). Monomeric, soluble forms of the ectodomain of the ACE2 and the Spike RBD-domain were expressed in human cells, to retain native glycosylation, and purified (Fig. S2). ACE2 was captured onto the sensor surface via a carboxy-terminal biotin and RBD injected over the ACE2 at different concentrations (Fig. 2A). Excellent fits of 1:1 Langmuir binding model to the data yielded an association rate constant (k_on_) of 0.9 ± 0.05 μM^−1^.s^−1^ and a dissociation rate constant (k_off_) of 0.067 ± 0.0011 s^−1^ (mean ± SD, n=6, Table 1). These rate constants are 3 to 25 fold faster than previously reported for the same interaction (Lei et al., 2020; Shang et al., 2020; Supasa et al., 2021; Wrapp et al., 2020; Zhang et al., 2021). However, previous experiments were conducted at unphysiologically low temperatures (i.e. below 37° C) and under conditions in which mass-transport limitations and rebinding are highly likely (see Discussion). These factors, and the presence of protein aggregates, would all lower the measured rate constants. In contrast, our measurements were conducted at 37° C and under conditions in which mass-transfer limitation and rebinding were excluded. The latter is demonstrated by the fact that measured k_on_ and k_off_ rates were clearly maximal at the low level of ACE2 immobilization (~50 RU) used in our experiments (Fig. 2B and C). The excellent fit of the 1:1 binding model to our data excludes an effect of protein aggregates, which yield complex kinetics. The calculated dissociation constant (K_D_) was 74 ± 4 nM (mean ± SD, n=6, Table 1). We also measured K_D_ by equilibrium binding (Fig. 2D), which avoids any artefacts induced by mass transfer limitations and rebinding. This K_D_ determined by equilibrium binding was very similar to the value calculated from kinetic data [63 ± 7.7 nM (mean ± SD, n = 24, Table 1], and did not vary with immobilization level (Fig. 2E), further validating our kinetic measurements. These affinity values are within the wide range reported in previous studies, which varied from K_D_ 11 to 133 nM (Lei et al., 2020; Shang et al., 2020; Supasa et al., 2021; Wrapp et al., 2020; Zhang et al., 2021).

**Table 1.** Affinity and kinetic data for RBD variants and ACE2 variants. Mean and SD of the k_off_, k_on_, calculated K_D_, and equilibrium K_D_ values for all RBD variants binding all ACE2 variants. For most measurements n = 3; the exceptions were RBD WT/ACE2 WT equilibrium K_D_ measurements (n =24) and other RBD WT measurements (n = 6). UK1, UK2, BR, SA refer to the B.1.1.7, VOC-202102-02, P2, and B.1.351 variants, respectively.

| | $k_{\text{off}}$ ($\text{s}^{-1}$) | SD | $k_{\text{on}}$ ($\mu\text{M}^{-1} \text{s}^{-1}$) | SD | $K_D$ Calc. (nM) | SD | $K_D$ Equi. (nM) | SD |
| --- | --- | --- | --- | --- | --- | --- | --- | --- |
| **RBD over WT ACE2** |  |  |  |  |  |  |  |  |
| WT | 0.0668 | 0.00113 | 0.90 | 0.05 | 74.4 | 4.0 | 62.6 | 7.7 |
| K417N | 0.177 | 0.00416 | 0.49 | 0.05 | 364 | 29 | 349 | 10 |
| K417T | 0.126 | 0.00510 | 0.55 | 0.04 | 230 | 23 | 226 | 19 |
| S477N | 0.0348 | 0.00037 | 0.81 | 0.03 | 42.9 | 2.1 | 42.6 | 3.0 |
| E484K | 0.0818 | 0.00183 | 1.54 | 0.03 | 53.1 | 1.7 | 52.6 | 2.0 |
| N501Y (UK1) | 0.0111 | 0.00017 | 1.59 | 0.04 | 7.0 | 0.25 | 5.5 | 2.4 |
| K417N/E484K | 0.251 | 0.00799 | 1.02 | 0.07 | 247 | 23 | 251 | 23 |
| K417T/E484K | 0.168 | 0.00573 | 1.10 | 0.05 | 153 | 12 | 147 | 8.6 |
| E484K/N501Y (UK2) | 0.0118 | 0.00037 | 2.33 | 0.10 | 5.1 | 0.36 | 3.7 | 2.7 |
| K417N/E484K/N501Y (SA) | 0.0291 | 0.00076 | 1.46 | 0.06 | 20.0 | 0.70 | 17.4 | 3.1 |
| K417T/E484K/N501Y (BR) | 0.0211 | 0.00021 | 1.56 | 0.07 | 13.5 | 0.45 | 12.2 | 3.4 |
| **RBD over S19P ACE2** |  |  |  |  |  |  |  |  |
| WT | 0.0298 | 0.00039 | 1.50 | 0.12 | 20.0 | 1.3 | 30.5 | 2.2 |
| K417N | 0.0782 | 0.00284 | 0.72 | 0.04 | 108 | 2.8 | 129 | 8.2 |
| K417T | 0.0521 | 0.00196 | 0.69 | 0.02 | 75.8 | 4.7 | 87.8 | 7.0 |
| S477N | 0.0257 | 0.00016 | 1.05 | 0.07 | 24.6 | 1.7 | 30.3 | 2.7 |
| E484K | 0.0325 | 0.00031 | 2.02 | 0.08 | 16.2 | 0.55 | 20.8 | 1.3 |
| N501Y (UK1) | 0.0051 | 0.00004 | 2.31 | 0.09 | 2.2 | 0.09 | 3.5 | 0.4 |
| K417N/E484K | 0.0961 | 0.00198 | 1.28 | 0.11 | 75.6 | 7.1 | 91.3 | 6.5 |
| K417T/E484K | 0.0660 | 0.00255 | 1.45 | 0.03 | 45.5 | 2.5 | 53.8 | 1.5 |
| E484K/N501Y (UK2) | 0.0051 | 0.00008 | 3.10 | 0.10 | 1.7 | 0.05 | 3.4 | 0.4 |
| K417N/E484K/N501Y (SA) | 0.0122 | 0.00009 | 2.16 | 0.03 | 5.7 | 0.07 | 10.4 | 1.2 |
| K417T/E484K/N501Y (BR) | 0.0085 | 0.00007 | 2.11 | 0.05 | 4.0 | 0.07 | 6.1 | 1.3 |
| **RBD over K26R ACE2** |  |  |  |  |  |  |  |  |
| S477N | 0.0240 | 0.00009 | 1.07 | 0.05 | 22.6 | 1.1 | 33.4 | 1.3 |
| WT | 0.0500 | 0.00062 | 1.60 | 0.16 | 31.4 | 2.6 | 48.8 | 2.5 |
| K417N | 0.154 | 0.00789 | 0.88 | 0.07 | 175 | 8.1 | 237 | 15 |
| K417T | 0.101 | 0.00079 | 0.81 | 0.12 | 127 | 17.4 | 154 | 2.8 |
| S477N | 0.0240 | 0.00009 | 1.07 | 0.05 | 22.6 | 1.1 | 33.4 | 1.3 |
| E484K | 0.0587 | 0.00109 | 2.03 | 0.03 | 28.9 | 1.0 | 35.9 | 1.5 |
| N501Y (UK1) | 0.0081 | 0.00002 | 2.34 | 0.09 | 3.5 | 0.15 | 7.5 | 1.5 |
| K417N/E484K | 0.191 | 0.00481 | 1.48 | 0.15 | 130 | 9.4 | 166 | 11 |
| K417T/E484K | 0.135 | 0.00407 | 1.53 | 0.02 | 88.0 | 3.9 | 105 | 0.7 |
| E484K/N501Y (UK2) | 0.0085 | 0.00018 | 3.06 | 0.23 | 2.8 | 0.17 | 6.4 | 0.3 |
| K417N/E484K/N501Y (SA) | 0.0234 | 0.00040 | 2.13 | 0.05 | 11.0 | 0.28 | 18.7 | 2.0 |
| K417T/E484K/N501Y (BR) | 0.0164 | 0.00028 | 2.21 | 0.06 | 7.4 | 0.33 | 15.3 | 0.8 |

**Figure 2.**
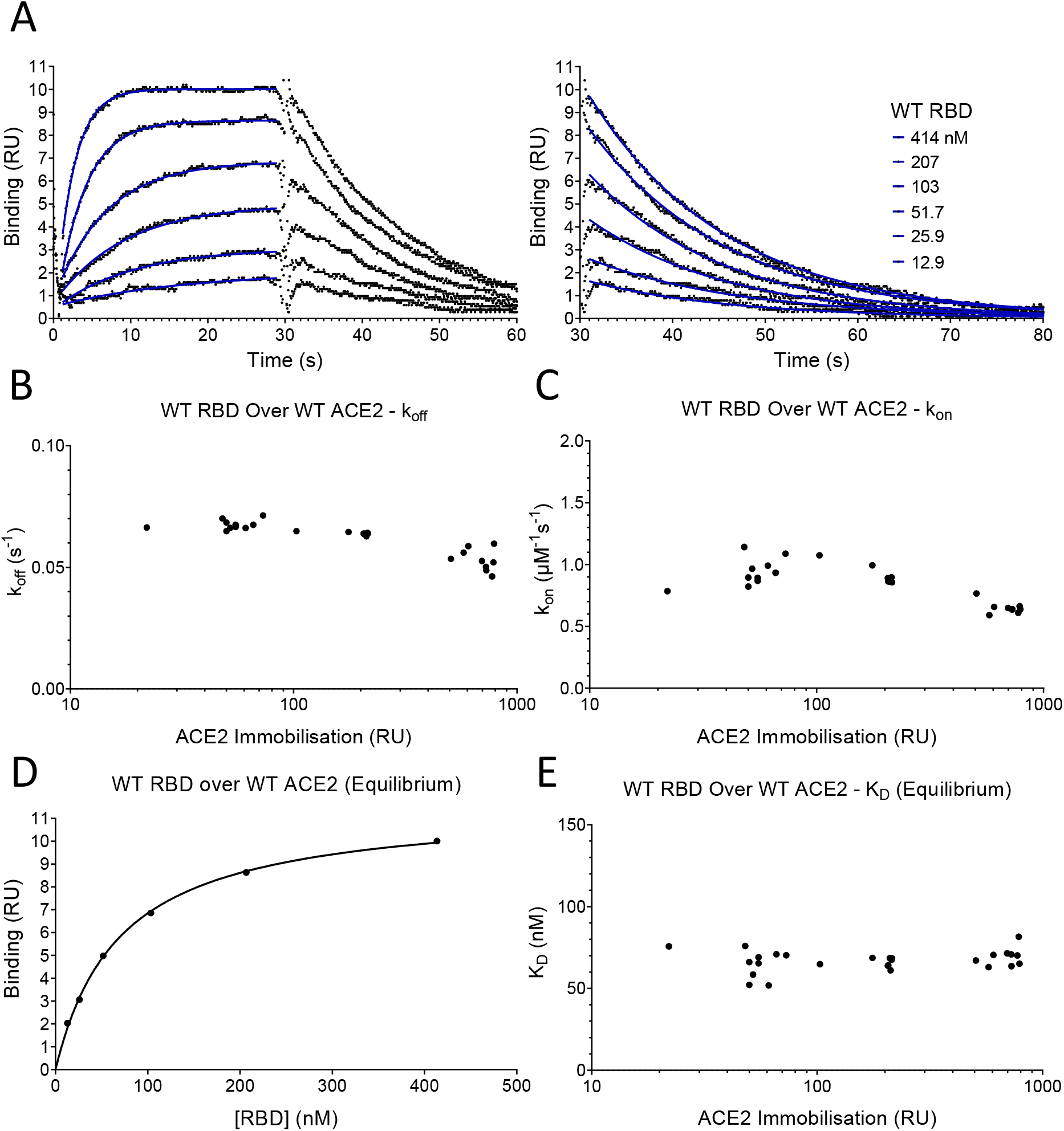
SPR analysis. (A) Overlay of binding traces showing association and dissociation when WT RBD is injected for 30 s at the indicated concentration over immobilized WT ACE2. The right panel shows an expanded view of the dissociation phase. The blue lines show the fits used for determining the k_on_ and k_off_. The k_on_ was determined as described in Fig. S3. The k_off_ (B) and k_on_ (C) values measured at different levels of immobilized ACE2 are shown. (D) The equilibrium K_D_ was determined by plotting the binding at equilibrium against [RBD] injected. Data from experiment shown in A. (E) The equilibrium K_D_ measured at different levels of immobilized ACE2 are shown.

### The effect of RBD mutations

We next evaluated the effect of RBD mutations on the affinity and kinetics of binding to ACE2 (Figure 3 and Table 1). Example sensorgrams are shown of mutations that increased (N501Y, Fig. 3A) or decreased (K417N, Fig. 3B) the binding affinity, while the key results from all mutants are summarized in Figure 3C. The single mutations S477N, E484K and N501Y all enhanced binding. The N501Y mutation had the biggest effect, increasing the affinity ~10 fold to K_D_ ~7 nM, by increasing the k_on_ ~1.8 fold and decreasing the k_off_ by ~ 7-fold. The S477N and E484K mutations increased the affinity more modestly (~ 1.5-fold), by decreasing the k_off_ (S477N) or increasing the k_on_ (E484K). The K417T and K417N mutations decreased the affinity ~2 and ~4 fold, respectively, mainly by decreasing the k_on_ but also by increasing the k_off_. Affinity-altering mutations in binding sites mainly affect the k_off_ (Agius et al., 2013) and have more modest effects on the k_on_. Changes in electrostatic interactions can dramatically affect the k_on_ (Schreiber and Fersht, 1996), and are a plausible explanation for the effects of the mutations K417T, K417N and E484K on k_on_. K417 forms a salt bridge with D30 on ACE2 (Lan et al., 2020) while E484 is ~9 Å from E75 on ACE2 (Lan et al., 2020). Thus the mutations K417N/T and E484K would decrease and increase, respectively, long-range electrostatic forces that may accelerate association (Schreiber and Fersht, 1996).

**Figure 3.**
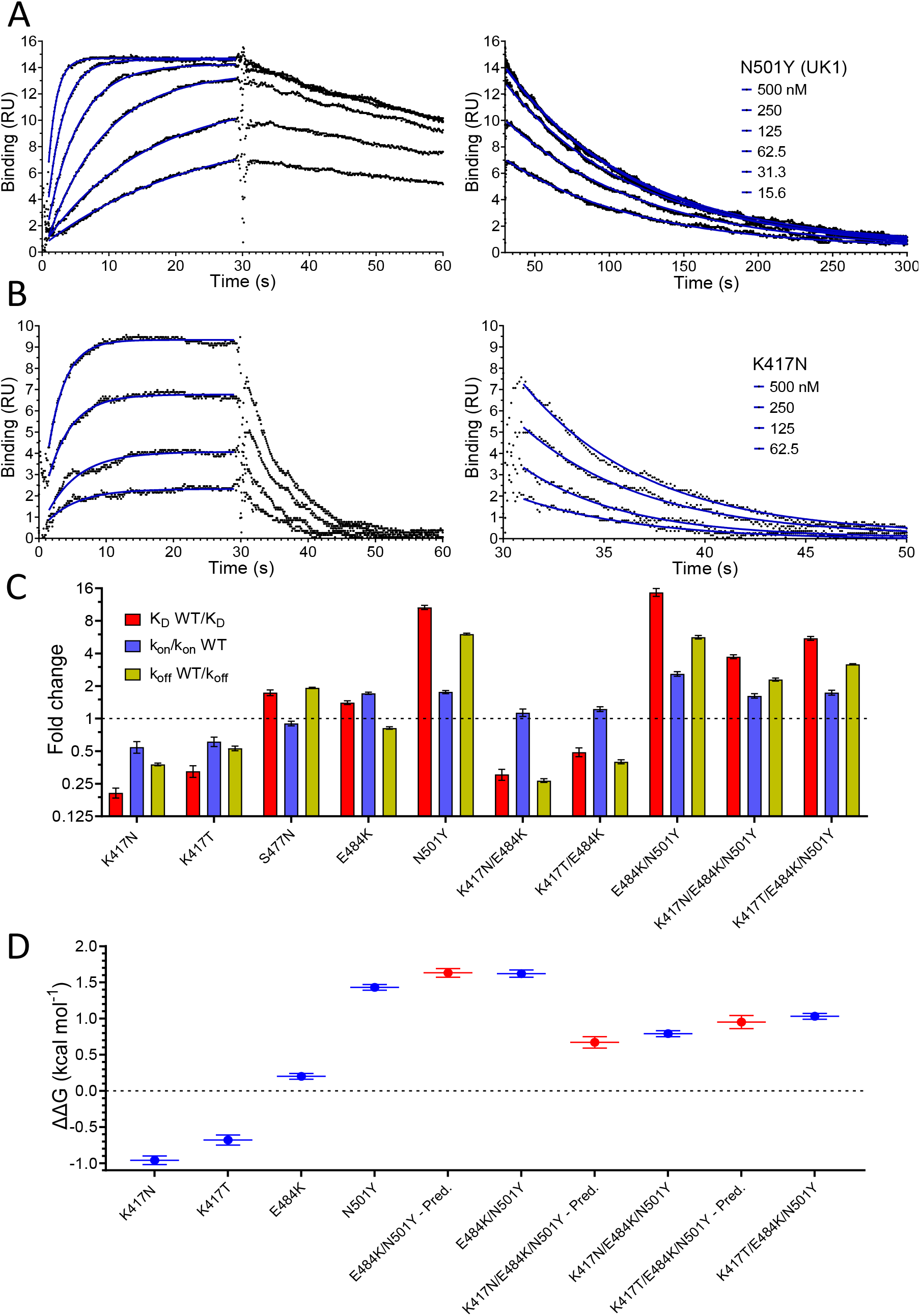
Effect of RBD variants binding WT ACE2. Overlay of binding traces showing association and dissociation of N501Y (A) and K417N (B) RBD variants when injected at a range of concentrations over immobilised WT ACE2. The right panels show an expanded view of the dissociation phase. The blue lines show fits used for determining the k_on_ and k_off_. (C) The fold change relative to WT RBD of the calculated K_D,_ k_on_, and k_off_ for the indicated RBD variants binding to immobilised WT ACE2 (Error bars show SD, n = 3). Representative sensorgrams from all mutants shown in Fig. S5, and the mean values from multiple repeats are in Table 1. (D) The blue lines show the measured ΔΔG for indicated RBD variants. The red lines show the predicted ΔΔG for the RBD variants with multiple mutations, which were calculated by adding ΔΔG values for single mutation variants (Error bars show SD, n = 3).

We also examined the effect on ACE2 binding of combinations of RBD mutations, including combinations present in VOC-202102-02, a subset of the B.1.1.7 lineage (N501Y) with the E484K mutation(“SARS-CoV-2 Variants of concern and variants under investigation - GOV.UK,” 2021), and the B.1.351 and P.1 variants (Fig. 3C, Table 1). In the case of VOC-202102-02, the addition of the E484K mutation to N501Y further increased the affinity, to ~15 fold higher than WT RBD (K_D_ ~5 nM), by further increasing the k_on_. Because the higher k_on_ could result in mass transfer limiting binding, we confirmed that the kinetic measurement for this variant was not substantially affected by varying levels of immobilization (Fig. S4). The affinity of the B.1.351 (K417N/ E484K/N501Y) and P.1 (K417T/E484K/N501Y) RBD variants for ACE2 increased by 3.7 and 5.3 fold, respectively, relative to wild type RBD, by both increasing the k_on_ and decreasing the k_off_ rate constants.

We next examined whether the effects of the mutations were additive, as is typically the case for multiple mutations at protein/protein interfaces (Wells, 1990). To do this we converted the changes in K_D_ to changes in binding energy (ΔΔG, Table 2) and examined whether the ΔΔG measured for RBD variants with multiple mutations was equal to the sum of the ΔΔG values measured for the individual RBD mutants. This was indeed the case (Fig. 3D), indicating that the effects on each mutation are independent. This is consistent with them being spaced well apart within the interface (Fig. 1C), and validates the accuracy of the affinity measurements.

**Table 2.** ΔΔG for RBD variants binding to ACE2 variants. Mean and SD of ΔΔG (n = 3, kcal/mol) were determined as described in the Materials and Methods using the calculated K_D_ values in Table 1. UK1, UK2, BR, and SA refer to the B.1.1.7, VOC-202102-02, P2, and B.1.351 variants, respectively.

|  | ACE2 WT |  | ACE2 S19P |  | ACE2 K26R |  |
| --- | --- | --- | --- | --- | --- | --- |
| RBD variant | $\Delta\Delta G$ | SD | $\Delta\Delta G$ | SD | $\Delta\Delta G$ | SD |
| WT | 0.00 | 0.00 | 0.79 | 0.05 | 0.52 | 0.06 |
| K417N | -0.96 | 0.06 | -0.23 | 0.04 | -0.52 | 0.04 |
| K417T | -0.68 | 0.07 | -0.01 | 0.05 | -0.32 | 0.09 |
| S477N | 0.33 | 0.04 | 0.67 | 0.05 | 0.72 | 0.04 |
| E484K | 0.20 | 0.04 | 0.92 | 0.04 | 0.57 | 0.04 |
| N501Y (UK1) | 1.43 | 0.04 | 2.13 | 0.04 | 1.86 | 0.04 |
| K417N/E484K | -0.72 | 0.07 | -0.01 | 0.07 | -0.34 | 0.06 |
| K417T/E484K | -0.43 | 0.06 | 0.30 | 0.05 | -0.10 | 0.04 |
| E484K/N501Y (UK2) | 1.62 | 0.05 | 2.30 | 0.04 | 1.98 | 0.05 |
| K417N/E484K/N501Y (SA) | 0.79 | 0.04 | 1.56 | 0.03 | 1.16 | 0.04 |
| K417T/E484K/N501Y (BR) | 1.03 | 0.04 | 1.76 | 0.03 | 1.39 | 0.04 |

### The effects of ACE2 mutations

We next examined the effects of mutations of ACE2 (S19P and K26R) on binding to both wild type and common variants of RBD (Fig. 4 and Table 1). Both S19P and K26R increased the affinity of WT RBD binding by ~3.7 and ~2.4 fold (Fig. 4A). These increases in affinity were the result of both increases in the k_on_ and decreases in the k_off_.

**Figure 4.**
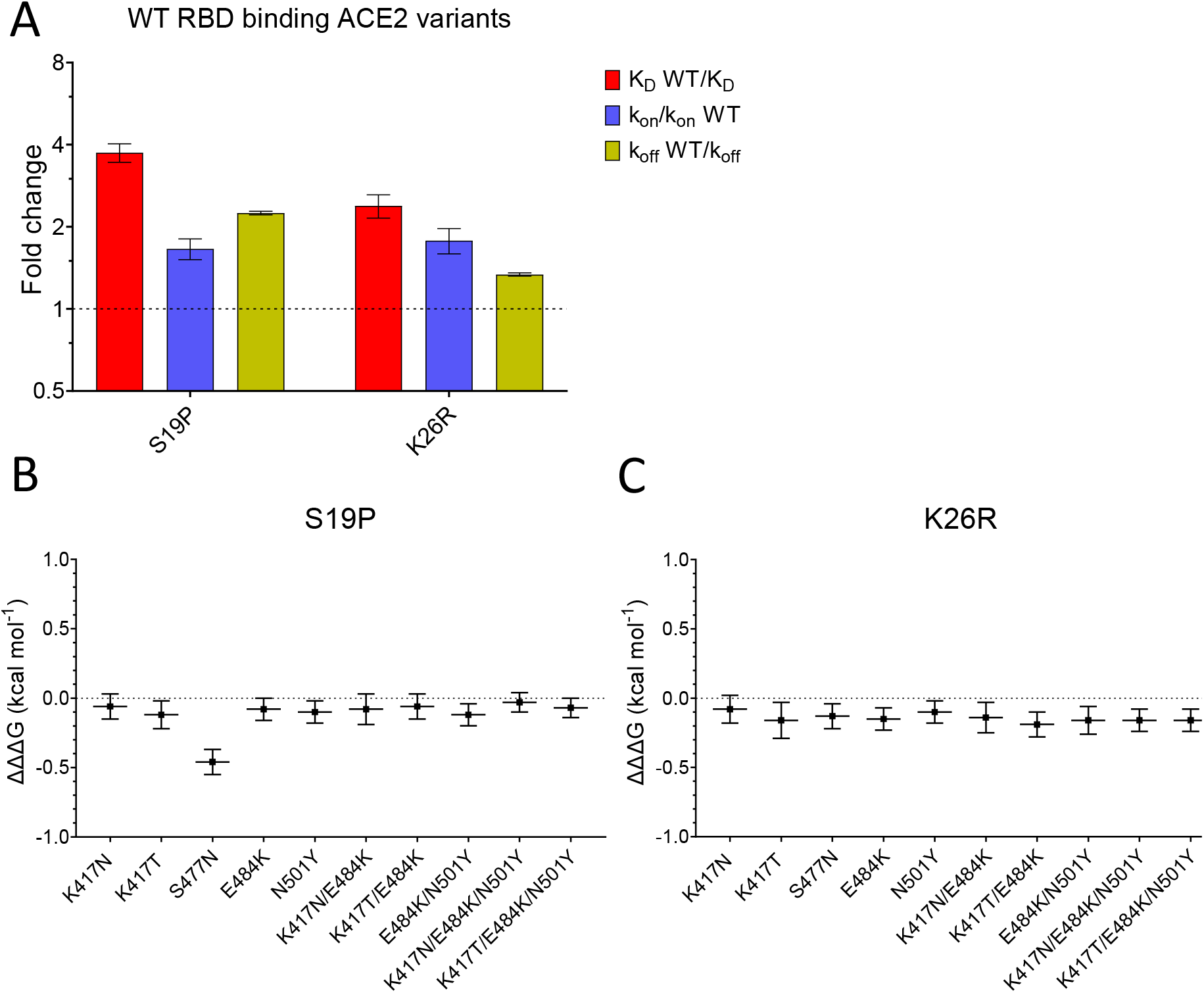
Effect of mutations in ACE2. (A) The fold change relative to WT ACE2 of the calculated K_D,_ k_on_, and k_off_ for the interaction of WT RBD and the indicated ACE2 variants (Error bars show SD, n = 3). (B-C) Show the difference (ΔΔΔG) between the measured and predicted ΔΔG for S19P (B) and K26R (C) ACE2 variants binding to the indicated RBD variants, calculated from data in Table 2. The predicted ΔΔG values for each variant RBD/variant ACE2 interaction were calculated from the sum of the ΔΔG for the ACE2 variant binding WT RBD and the ΔΔG for the RBD variant binding WT ACE2 (Table 2).

Finally, we looked for interactions between RBD and ACE2 mutations by measuring the effects of the ACE2 mutations on binding to all mutant forms of RBD (Table 1). After converting changes in K_D_ to ΔΔG (Table 2) we examined whether ΔΔG measured for a given ACE2 variant/RBD variant interaction was equal to the sum of the ΔΔG measured for ACE2 variant/RBD WT and ACE WT/RBD variant interactions. This is depicted as the difference between the measured and predicted ΔΔG for interactions between ACE2 and RBD variants (ΔΔΔG in Figs. 4B and C). In most cases ΔΔΔG values were close to zero, indicating that the effects of these mutations were largely independent. The one exception was the combination of ACE2 S19P and RBD S477N variants, where the measured value was significantly lower than the predicted value (Fig. 4B), indicating that these mutations were not independent. This is consistent with the fact that the ACE2 residue S19 is adjacent to RBD residue S477 in the contact interface (Fig. 1C). An important consequence of this is that the S477N mutation increased the affinity of RBD for ACE2 WT but decreased its affinity for ACE2 S19P.

## Discussion

While our finding that the SARS-CoV-2 RBD binds ACE2 with an affinity of K_D_ 74 nM at 37°C is consistent with previous studies (K_D_ 11 to 133 nM) (Lei et al., 2020; Shang et al., 2020; Supasa et al., 2021; Wrapp et al., 2020; Zhang et al., 2021, 2020), the rate constants that we measured (k_on_ 0.9 μM^−1^.s^−1^ and k_off_ 0.067 s^−1^) were more than 3 fold faster than all previous reports. One likely reason for this is that previous measurements were performed at a lower temperature, which almost always decreases rate constants. While one study stated that binding constants were measured at 25°C (Zhang et al., 2020), most studies did not report the temperature, suggesting that they were performed at room temperature or the standard instrument temperature (20-25°C). A second likely reason is that previous kinetic studies were performed under conditions in which the rate of diffusion of soluble molecule to the sensor surface limits the association rate, and rebinding of dissociated molecules to the surface reduces the measured dissociation rate. These are known pitfalls of both techniques used in these studies, surface plasmon resonance (Myszka, 1997) and bilayer interferometry (Abdiche et al., 2008). In the present study we avoided these issues by immobilizing a very low level of ligand on the sensor surface. A third possible reason is that the proteins were aggregated, which can cause problems even when aggregates are a very minor contaminant (van der Merwe and Barclay, 1996). The presence of aggregates results in complex binding kinetics, which can be excluded if the simple 1:1 Langmuir binding model fits the kinetic data. While this was demonstrated in the present study, and some previous studies (Shang et al., 2020; Wrapp et al., 2020; Zhang et al., 2021), such fits were not shown in all studies, one of which reported more than 20 fold slower kinetics than reported here (Lei et al., 2020; Supasa et al., 2021).

The RBD mutants that we selected for analysis have all emerged independently and become dominant in a region at least once in different lineages, suggesting that they provide a selective advantage. Our finding that the N501Y, E484K, and S477N all increase the binding affinity of RBD for ACE2 raises the question as to whether this contributed to their selection. Several lines of evidence suggest that enhancing the Spike/ACE2 interaction would be advantageous. Firstly, the virus has spread only very recently to humans from another mammalian host, providing insufficient time for optimization of the affinity. Secondly, epidemiological studies have demonstrated enhanced transmissibility of the B.1.1.7 variant, which has the N501Y mutation (Volz et al., 2021b; Washington et al., 2021). Finally, a SARS-CoV-2 variant with the Spike mutation D614G, which increases its activity by stabilizing it following furin cleavage (Zhang et al., 2021, 2020), rapidly became dominant globally after it emerged (Korber et al., 2020; Volz et al., 2021a). Taken together, these findings suggest that the WT Spike/ACE2 interaction is limiting for transmission, and that mutations which enhance it, including the N501Y, E484K, and S477N mutations, would provide a selective advantage by increasing transmissibility. This raises two questions. Firstly, will other RBD mutations appear in SARS-CoV-2 which further enhance transmission? This seems likely, given that a large number of RBD mutations have been identified that increase the RBD/ACE2 affinity (Starr et al., 2020; Zahradník et al., 2021). Secondly, will combinations of existing mutations be selected because they further increasing the affinity? While the appearance E484K together with the N501Y in three lineages (B.1.1.7, B.1.351 and P.1) supports this, it is also possible that E484K was selected because it disrupts antibody neutralization, as discussed below.

Studies of other enveloped viruses, including SARS-CoV-2, suggest that increases in affinity of viral fusion ligands for their cellular receptors can increase cell infection and disease severity (Hasegawa et al., 2007; Li et al., 2005). One study found that increasing this affinity enabled the virus to infect cells with lower receptor surface density (Hasegawa et al., 2007). It follows that increases in affinity could increase the number of host tissues infected, which could increase the severity of disease (Cao and Li, 2020) and/or increase the viral load in the upper respiratory tract el (Hoffmann et al., 2020; Wölfel et al., 2020), thereby increasing spread.

Another mechanism by which mutations of RBD could provide a selective advantage is through evasion of immune responses. This is supported by the observation that neutralizing antibodies present in those infected by or vaccinated against SARS-CoV-2 primarily target the RBD domain (Garcia-Beltran et al., 2021; Greaney et al., 2021a; Rogers et al., 2020). Furthermore, two variants with RBD mutations that abrogate antibody neutralization, B.1.351 and P1, became dominant in regions with very high levels of prior SARS-CoV-2 infection (Cele et al., 2021; Dejnirattisai et al., 2021; Hoffmann et al., 2021; Sabino et al., 2021; Tegally et al., 2021; D. Zhou et al., 2021). Both lineages include the N501Y mutation, but this appears to have modest effects on antibody neutralization (Greaney et al., 2021a, 2021b). In contrast, the E484K mutation, also present in both lineages, potently disrupts antibody neutralization (Greaney et al., 2021a, 2021b). Our finding that the K417N/T mutants present in B.1.351/P.1 lineages decrease the affinity of RBD for ACE2 suggests that they were selected because they facilitate immune escape. Indeed, mutations of K417 can block antibody neutralization, albeit less effectively than E484K (Greaney et al., 2021a, 2021b; Wang et al., 2021). It is notable that these affinity-reducing K417N/T mutants have only emerged together with mutants (N501Y and E484K) that increase the affinity of RBD for ACE2, suggesting a cooperative effect between mutations that enhance immune escape and mutations that increase affinity.

The effect of the increased affinity for SARS-CoV-2 Spike RBD of the K26R and S19P ACE2 mutants are less clear. The evidence summarised above that WT RBD/ACE2 binding is limiting for SARS-CoV-2 transmission, suggest that carriers of these ACE2 variants will be at greater risk of infection and/or severe disease. However, in contrast to SARS-CoV-2 RBD mutations, the effects of ACE2 variants are primarily relevant to the carriers of these mutations. A preliminary analysis (MacGowan et al., 2021) suggests that the carriers of the K26R ACE allele might be at increased risk of severe disease, but the findings did not reach statistical significance, and further studies are required.

The interaction that we identified between the RBD S477N and ACE2 S19P mutants highlights the importance of considering variation in the host population when studying the evolution of viral variants. In this case, the opposite effect of the RBD S477N mutation on its affinity for ACE2 S19P (decreased) compared with ACE2 WT (increased), suggests that this RBD variant may have a selective disadvantage amongst carriers of the ACE2 S19P variant, in contrast to those with ACE2 WT, where it appears to be advantageous. However, the low frequency of this variant means that this is unlikely to be important at a population level and will be difficult to detect.

It is noteworthy that the two most common ACE2 variants are in positions on ACE2 with no known functional activity. This raises the question as to whether these mutations are a remnant of historic adaption to pathogens that utilised this portion of ACE2. The fact that ACE2 S19P mutation is largely confined to African/African-American populations, suggests that it is more recent than K26R and/or selected by pathogen(s) confined to the African continent.

## Materials and Methods

### ACE2 and RBD variant constructs

The soluble WT ACE2 construct, which was kindly provided by Ray Owens (Oxford Protein Production Facility-UK), encoded the following protein:

STIEEQAKTFLDKFNHEAEDLFYQSSLASWNYNTNITEENVQNMNNAGDKWSAFLKEQSTLAQMYPLQEIQNLTVKLQLQALQQNGSSVLSEDKSKRLNTILNTMSTIYSTGKVCNPDNPQECLLLEPGLNEIMANSLDYNERLWAWESWRSEVGKQLRPLYEEYVVLKNEMARANHYEDYGDYWRGDYEVNGVDGYDYSRGQLIEDVEHTFEEIKPLYEHLHAYVRAKLMNAYPSYISPIGCLPAHLLGDMWGRFWTNLYSLTVPFGQKPNIDVTDAMVDQAWDAQRIFKEAEKFFVSVGLPNMTQGFWENSMLTDPGNVQKAVCHPTAWDLGKGDFRILMCTKVTMDDFLTAHHEMGHIQYDMAYAAQPFLLRNGANEGFHEAVGEIMSLSAATPKHLKSIGLLSPDFQEDNETEINFLLKQALTIVGTLPFTYMLEKWRWMVFKGEIPKDQWMKKWWEMKREIVGVVEPVPHDETYCDPASLFHVSNDYSFIRYYTRTLYQFQFQEALCQAAKHEGPLHKCDISNSTEAGQKLFNMLRLGK SEPWTLALENVVGAKNMNVRPLLNYFEPLFTWLKDQNKNSFVGWSTDWSPYAD<u>LNDIFEAQKIEWHE</u> <u>K</u>HHHHHH

The carboxy-terminal end has a biotin acceptor peptide (underlined) followed by an oligohistidine tag.

The WT RBD construct, which was kindly provided by Quentin Sattentau (Sir William Dunn School of Pathology), encoded the following protein:

RVQPTESIVRFPNITNLCPFGEVFNATRFASVYAWNRKRISNCVADYSVLYNSASFSTFKCYGVSPTKLNDLCFTNVYADSFVIRGDEVRQIAPGQTGKIADYNYKLPDDFTGCVIAWNSNNLDSKVGGNYNYLYRLFRKSNLKPFERDISTEIYQAGSTPCNGVEGFNCYFPLQSYGFQPTNGVGYQPYRVVVLSFELLHAPATVCGPKKSTNLVKNKCVNFHHHHHH

The carboxy-terminal end has an oligohistidine tag.

ACE2 and RBD point mutations were introduced using the Agilent QuikChange II XL Site-Directed Mutagenesis Kit following the manufacturer’s instructions. The primers were designed using the Agilent QuikChange primer design web program.

### HEK293F cell transfection

Cells were grown in FreeStyle™ 293 Expression Medium (12338018) in a 37 °C incubator with 8% CO_2_ on a shaking platform at 130 rpm. Cells were passaged every 2-3 days with the suspension volume always kept below 33.3% of the total flask capacity. The cell density was kept between 0.5 and 2 million per ml. Before transfection cells were counted to check cell viability was above 95% and the density adjusted to 1.0 million per ml. For 100 ml transfection, 100 μl FreeStyle™ MAX Reagent (16447100) was mixed with 2 ml Opti-MEM (51985034) for 5 minutes. During this incubation 100 μg of expression plasmid was mixed with 2 ml Opti-MEM. For in situ biotinylation of ACE2 90 μg of expression plasmid was mixed with 10 μg of expression plasmid encoding the BirA enzyme. The DNA was then mixed with the MAX Reagent and incubated for 25 minutes before being added to the cell culture. For ACE2 in situ biotinylation, biotin was added to the cell culture at a final concentration of 50 μM. The culture was left for 5 days for protein expression to take place.

### Protein purification

Cells were harvested by centrifugation and the supernatant collected and filtered through a 0.22 μm filter. Imidazole was added to a final concentration of 10 mM and PMSF added to a final concentration of 1 mM. 1 ml of Ni-NTA Agarose (30310) was added per 100 ml of supernatant and the mix was left on a rolling platform at 4 °C overnight. The supernatant mix was poured through a gravity flow column to collect the Ni-NTA Agarose. The Ni-NTA Agarose was washed 3 times with 25 ml of wash buffer (50 mM NaH2PO4, 300 mM NaCl and 20 mM imidazole at pH 8). The protein was eluted from the Ni-NTA Agarose with elution buffer (50 mM NaH_2_PO_4_, 300 mM NaCl and 250 mM imidazole at pH 8). The protein was concentrated, and buffer exchanged into size exclusion buffer (25 mM NaH_2_PO_4_, 150 mM NaCl at pH 7.5) using a protein concentrator with a 10,000 molecular weight cut-off. The protein was concentrated down to less than 500 μl before loading onto a Superdex 200 10/300 GL size exclusion column (Fig. S2). Fractions corresponding to the desired peak were pooled and frozen at -80 °C. Samples from all observed peaks were analysed on an SDS-PAGE gel (Fig. S2).

### Surface plasmon resonance (SPR)

RBD binding to ACE2 was analysed on a Biacore T200 instrument (GE Healthcare Life Sciences) at 37°C and a flow rate of 30 μl/min. Running buffer was HBS-EP (BR100669). Streptavidin was coupled to a CM5 sensor chip (29149603) using an amine coupling kit (BR100050) to near saturation, typically 10000-12000 response units (RU). Biotinylated ACE2 WT and variants were injected into the experimental flow cells (FC2–FC4) for different lengths of time to produce desired immobilisation levels (40–800 RU). FC1 was used as a reference and contained streptavidin only. Excess streptavidin was blocked with two 40 s injections of 250 μM biotin (Avidity). Before RBD injections, the chip surface was conditioned with 8 injections of the running buffer. A dilution series of RBD was then injected in all FCs. Buffer alone was injected after every 2 or 3 RBD injections. The length of all injections was 30 s, and dissociation was monitored from 180-670 s. The background response measured in FC1 was subtracted from the response in the other three FCs. In addition, the responses measured during buffer injections closest in time were subtracted. Such double-referencing improves data quality when binding responses are low as needed to obtain accurate kinetic data (Myszka, 1999). At the end of each experiment an ACE2-specific mouse monoclonal antibody (NOVUS Biologicals, AC384) was injected at 5 μg/ml for 10 minutes to confirm the presence and amount of immobilized ACE2.

### Data analysis

Double referenced binding data was fitted using GraphPad Prism. The k_off_ was determined by fitting a mono-exponential decay curve to data from the dissociation phase of each injection. The k_off_ from four to six RBD injections was averaged to give a value for the k_off_ (Fig. S3A). The k_on_ was determined by first fitting a mono-exponential association curve to data from the association phase, yielding the k_obs_. The k_on_ was be determined by plotting the k_obs_ vs the concentration of RBD and performing a linear fit of the equation k_obs_ = k_on_*[RBD] + k_off_ to this data (Fig. S3B), using the k_off_ determined as above to constrain the fit.

The K_D_ was either calculated (calculated K_D_ = k_off_/k_on_) or measured directly (equilibrium K_D_) as follows. Equilibrium binding levels at a given [RBD] were determined from the fit above of the mono-exponential association phase model to the association phase data. These equilibrium binding levels were plotted against [RBD] and a fit of the simple 1:1 Langmuir binding model to this data was used to determine the equilibrium K_D_ (Fig. 2D).

ΔG for each affinity measurement was calculated the relationship ΔG =R*T*lnK_D_, where R = 1.987 cal mol^−1^ K^−1^, T = 310.18 K, and K_D_ is in units M. ΔΔG values (Table 2 and Fig. 3D) were calculated for each mutant from the relationship ΔΔG = ΔG_WT_ — ΔG_M_. The predicted ΔΔG for interactions with multiple mutants were calculated by adding the single mutant ΔΔG values (Fig. 3D). The difference between the measured and predicted ΔΔG (ΔΔΔG) for interactions between the ACE2 and RBD mutants was calculates as ΔΔΔG = measured ΔΔG – predicted ΔΔG (Fig. 4B).

All errors represent standard deviations and errors for calculated values were determined by error propagation.

## Acknowledgments

We thank Johannes Pettmann for help with protein expression and Anna Huhn for help with data analysis. OD is supported by a Wellcome Trust Senior Fellowship in Basic Biomedical Sciences (207537/Z/17/Z828). SM and GB are supported by Biotechnology and Biological Sciences Research Council Grants (BB/J019364/1 and BB/R014752/1) and a Wellcome Trust Biomedical Resources Grant (101651/Z/13/Z).

---

**Figure S1.**
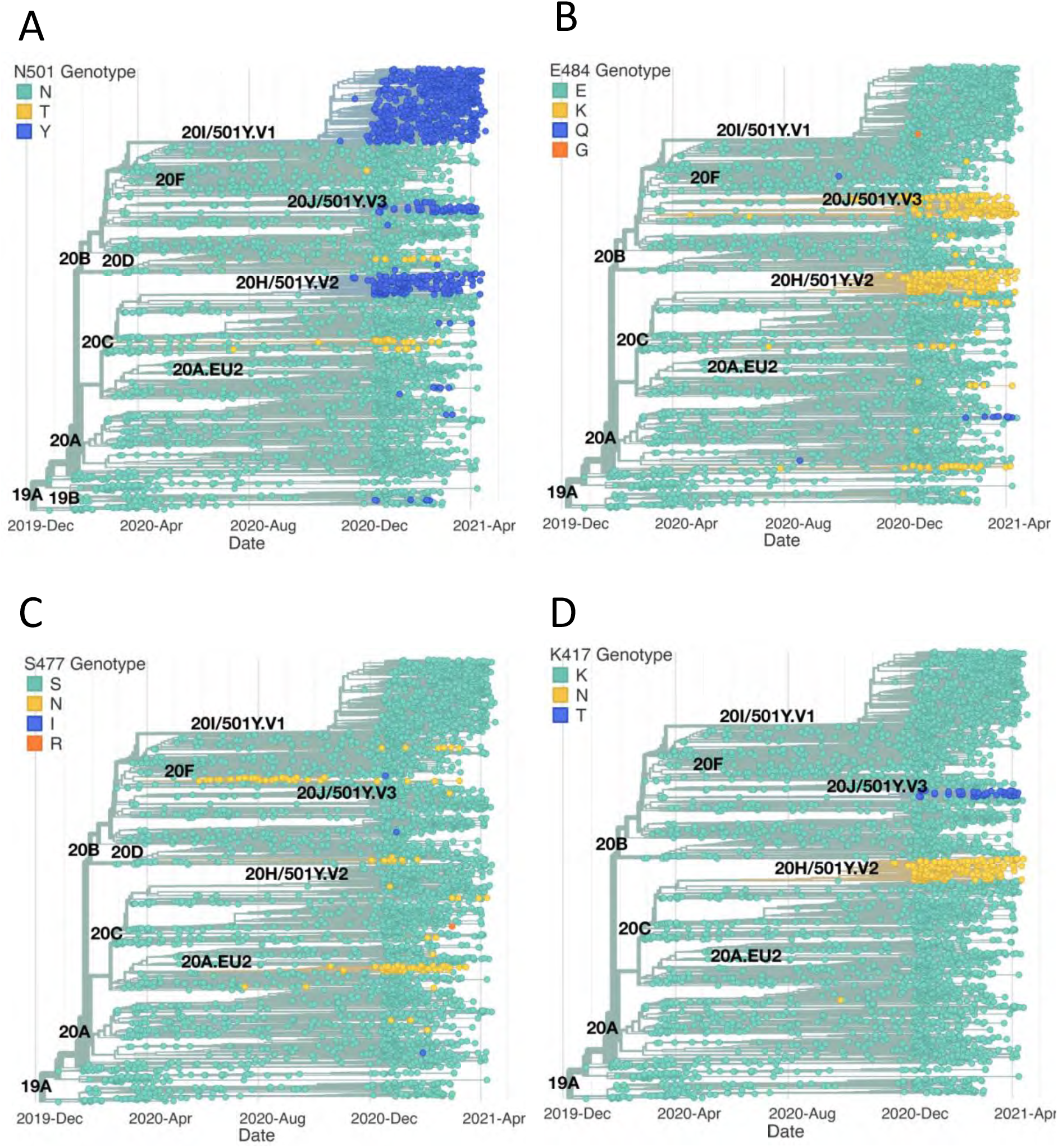
Emergence of the same RBD mutations in multiple SAR2-CoV-2 clades. The figure highlights the SARS-CoV-2 clades containing RBD mutations investigated in this study. The phylogenetic trees were constructed as in Fig. 1A from SARS-CoV-2 sequences accessed on the 22nd April 2021 (N = 3,914). (A) N501Y has emerged independently of the three clades 501Y.V1, 501Y.V2, and 501Y.V3. Mutation to T at this position has also occurred frequently. (B) E484K has also been observed independently of its main progenitor clades 501Y.V2 and 501Y.V3. E484Q and E484G have also been observed. (C) S477N has been observed beyond clades 20F and 20A.EU2. Mutations to I and R have also been occasionally observed at this position. (D) Mutations of K417 to N and T have been observed almost exclusively in the 20H.501Y.V2 and 20J.501Y.V3 clades.

**Figure S2.**
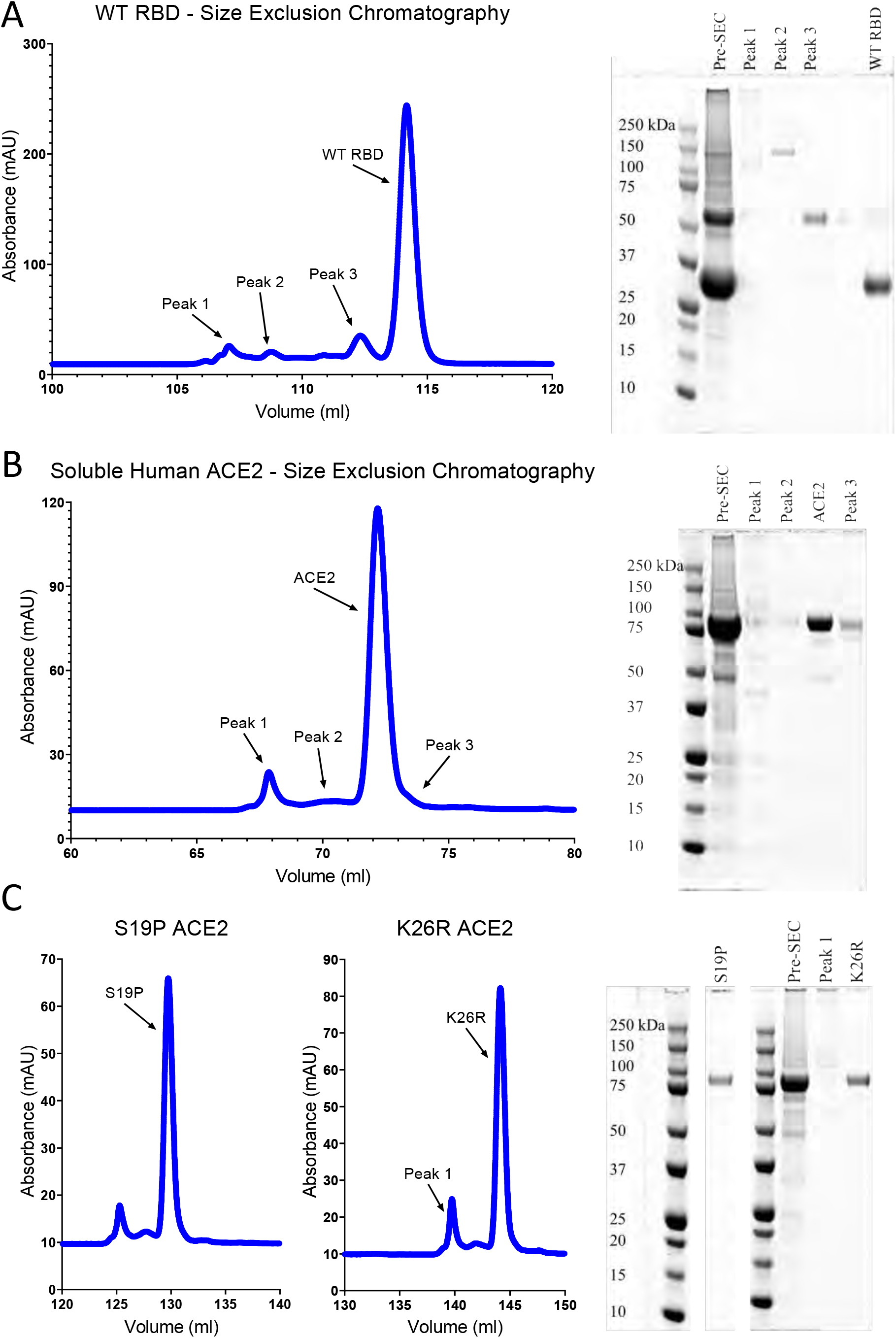
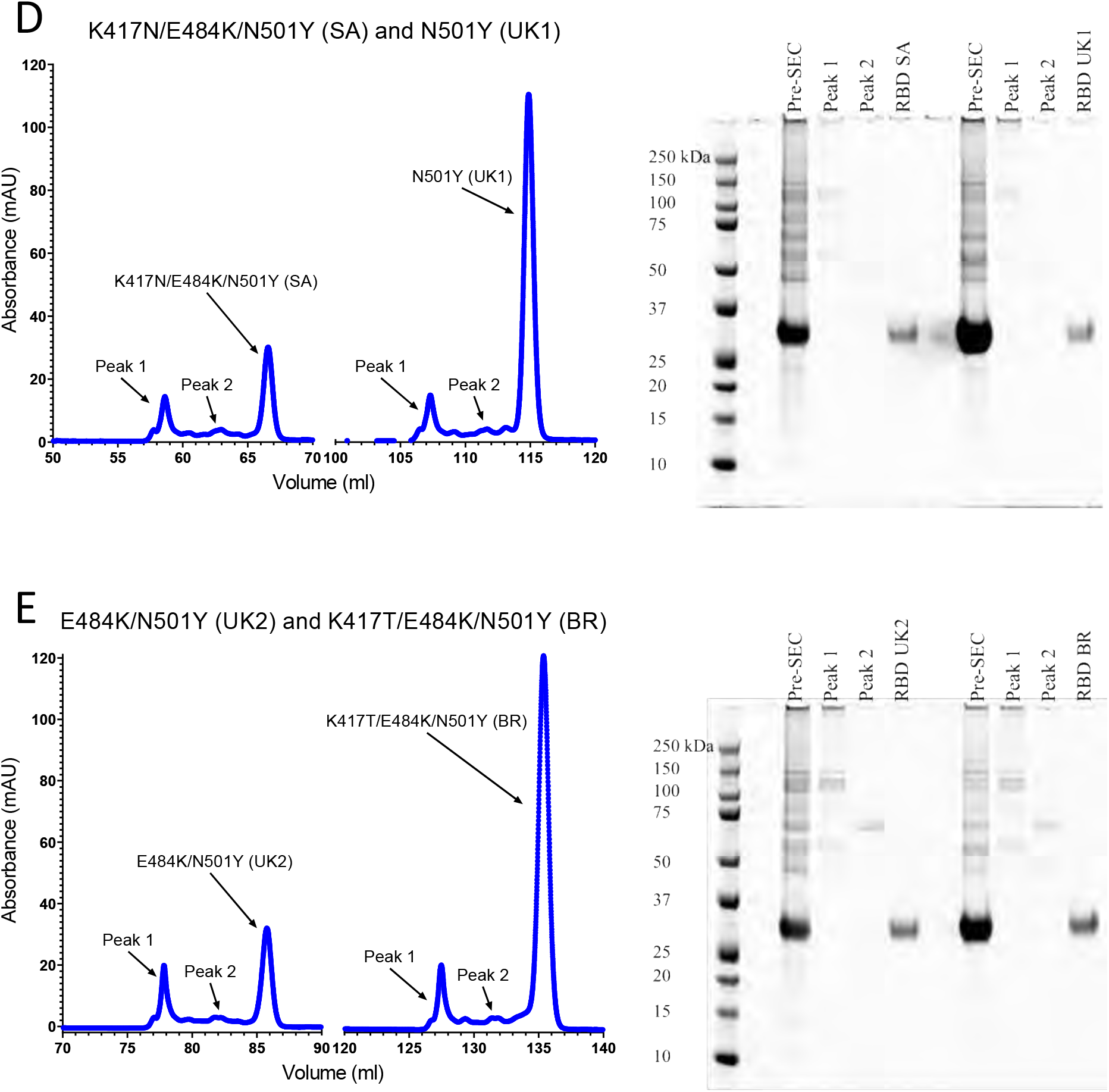
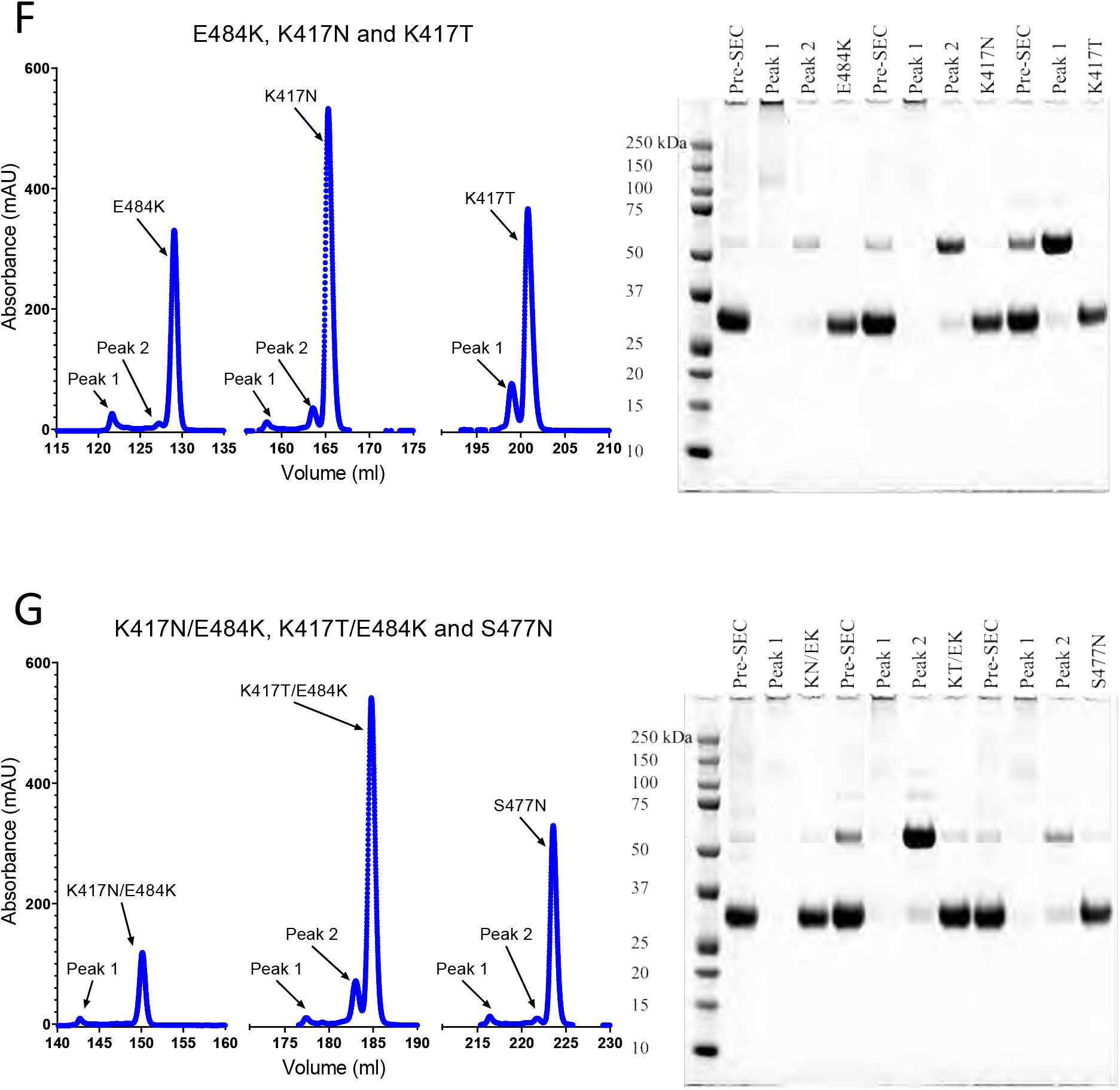
Protein purification. Size- exclusion chromatography traces of the indicated ACE2 and RBD proteins and SDS-PAGE of the indicated peak fractions. UK1, UK2, BR, SA refer to the B.1.1.7, VOC-202102-02, P2, and B.1.351 variants, respectively.

**Figure S3.**
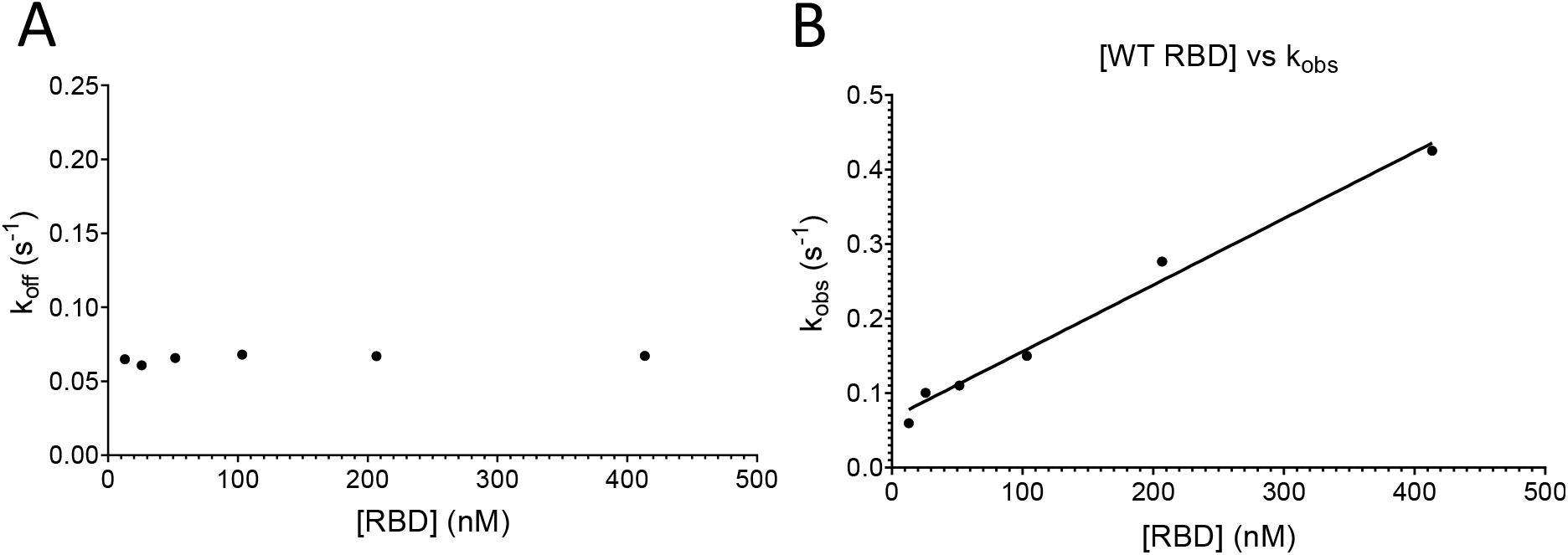
Determining the k_on_ and k_off_. Analysis of data from the fits in Fig. 2A. (A) A plot of k_off_ obtained for each injection versus [RBD]. (B) A plot of k_obs_ for each injection versus [RBD]. The line shows a constrained fit of the equation k_obs_ = k_on_*[RBD] + k_off_, using the k_off_ obtained in (A). The k_on_ was obtained from the slope.

**Figure S4.**
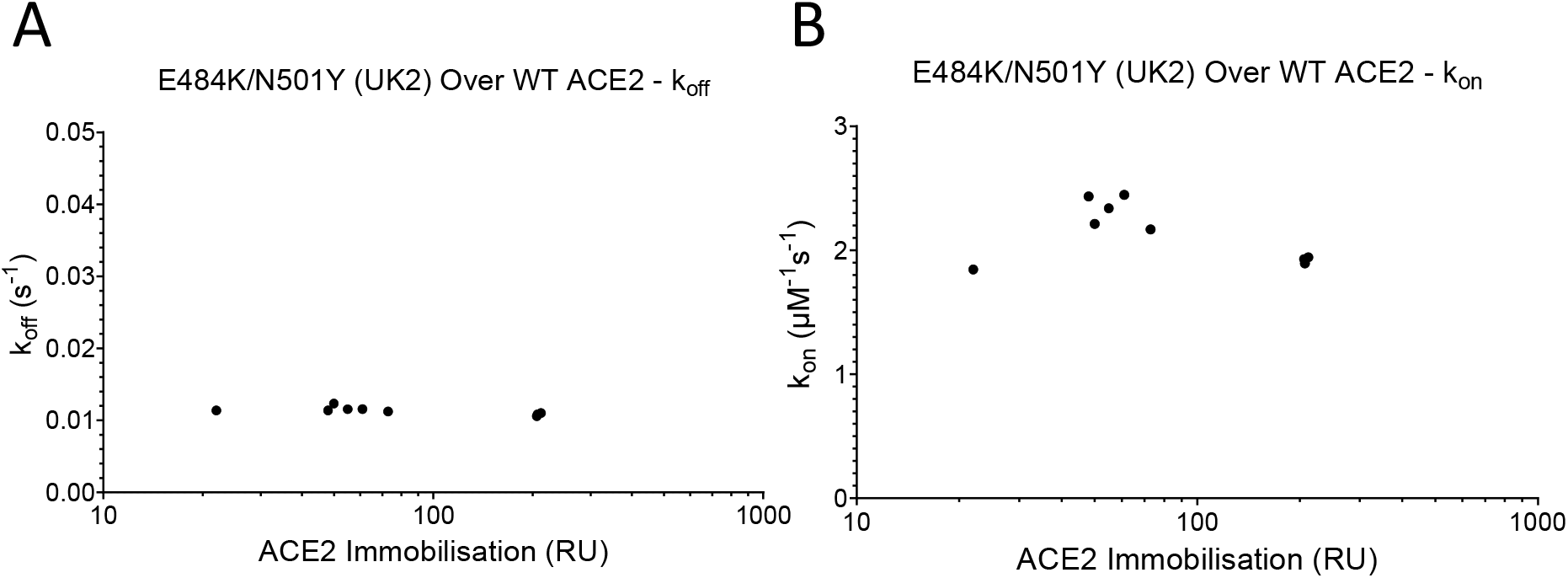
Mass transport controls from RBD. The k_off_ (A) and k_on_ (B), respectively, for E484K/N501Y (UK2) RBD binding WT ACE2 at a range of surface immobilisations (n = 12). UK2 refers to VOC-202102-02.

**Figure S5.**
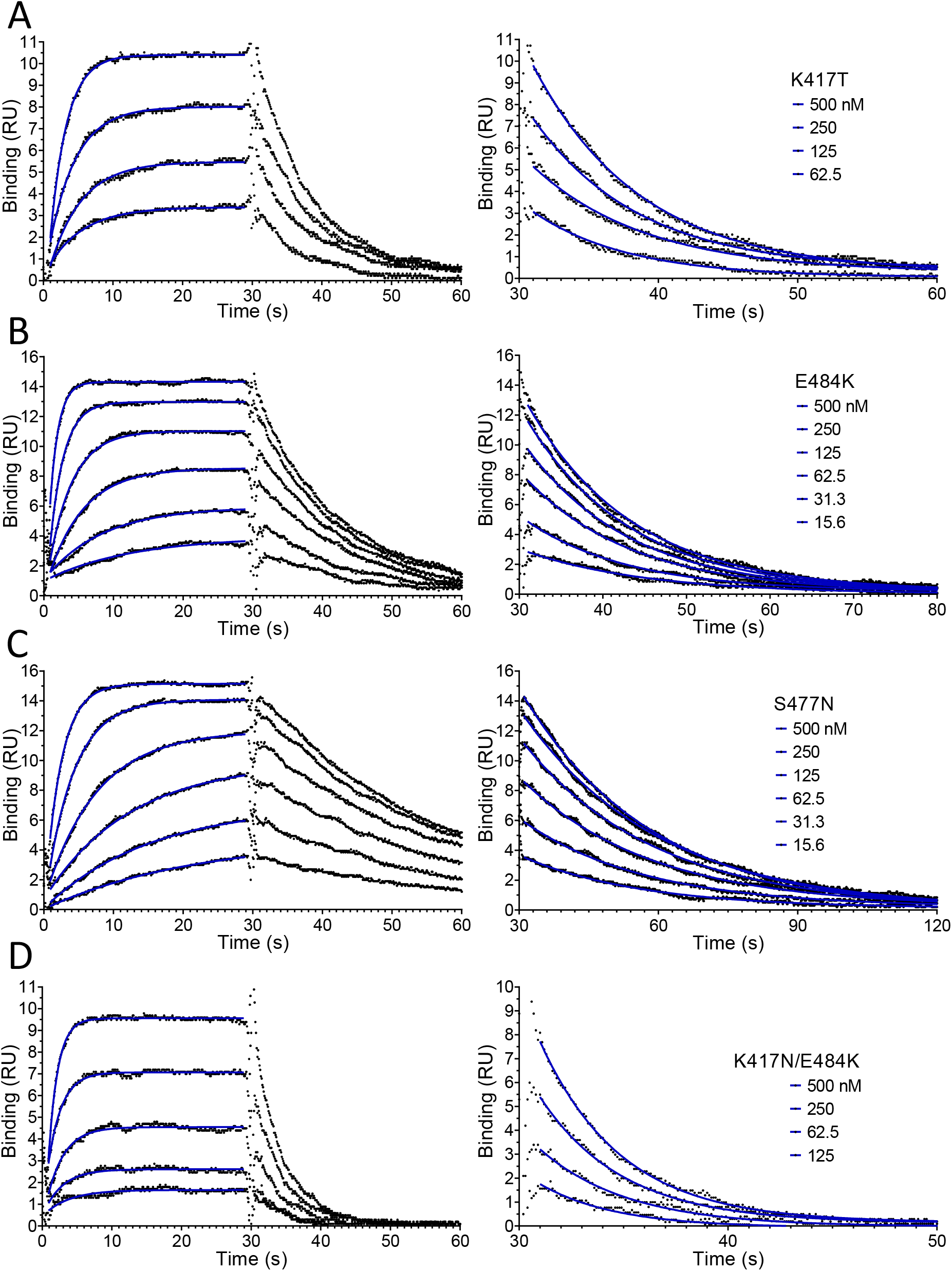
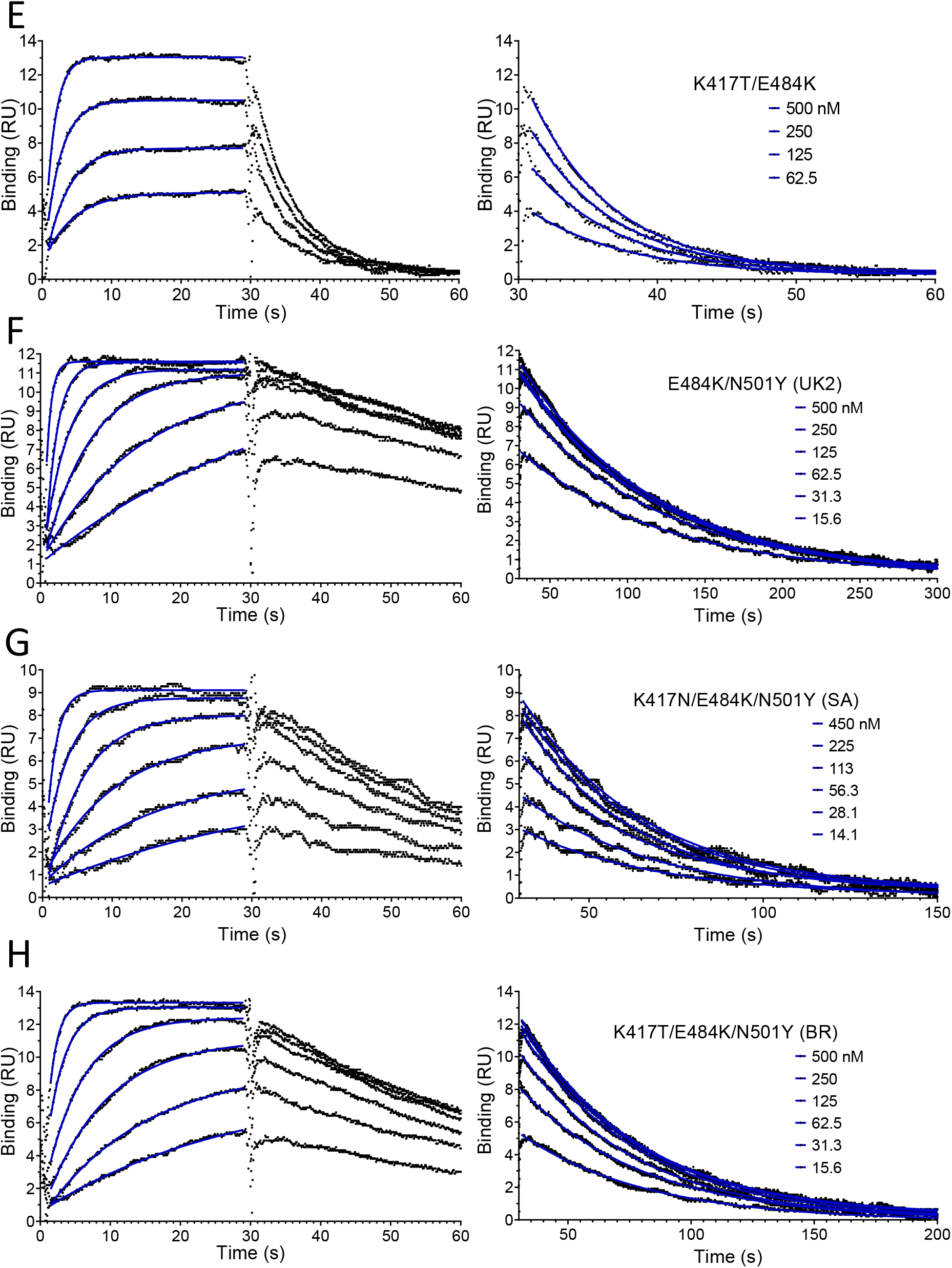
Representative SPR data for RBD variants binding to WT ACE2. Binding traces for the indicated RBD variants injected different concentrations over immobilised WT ACE2. The right panels show an expanded view of the dissociation phase. The blue lines show fits used for determining the k_on_ and k_off_. UK1, UK2, BR, SA refer to the B.1.1.7, VOC-202102-02, P2, and B.1.351 variants, respectively.

**Figure S6.**
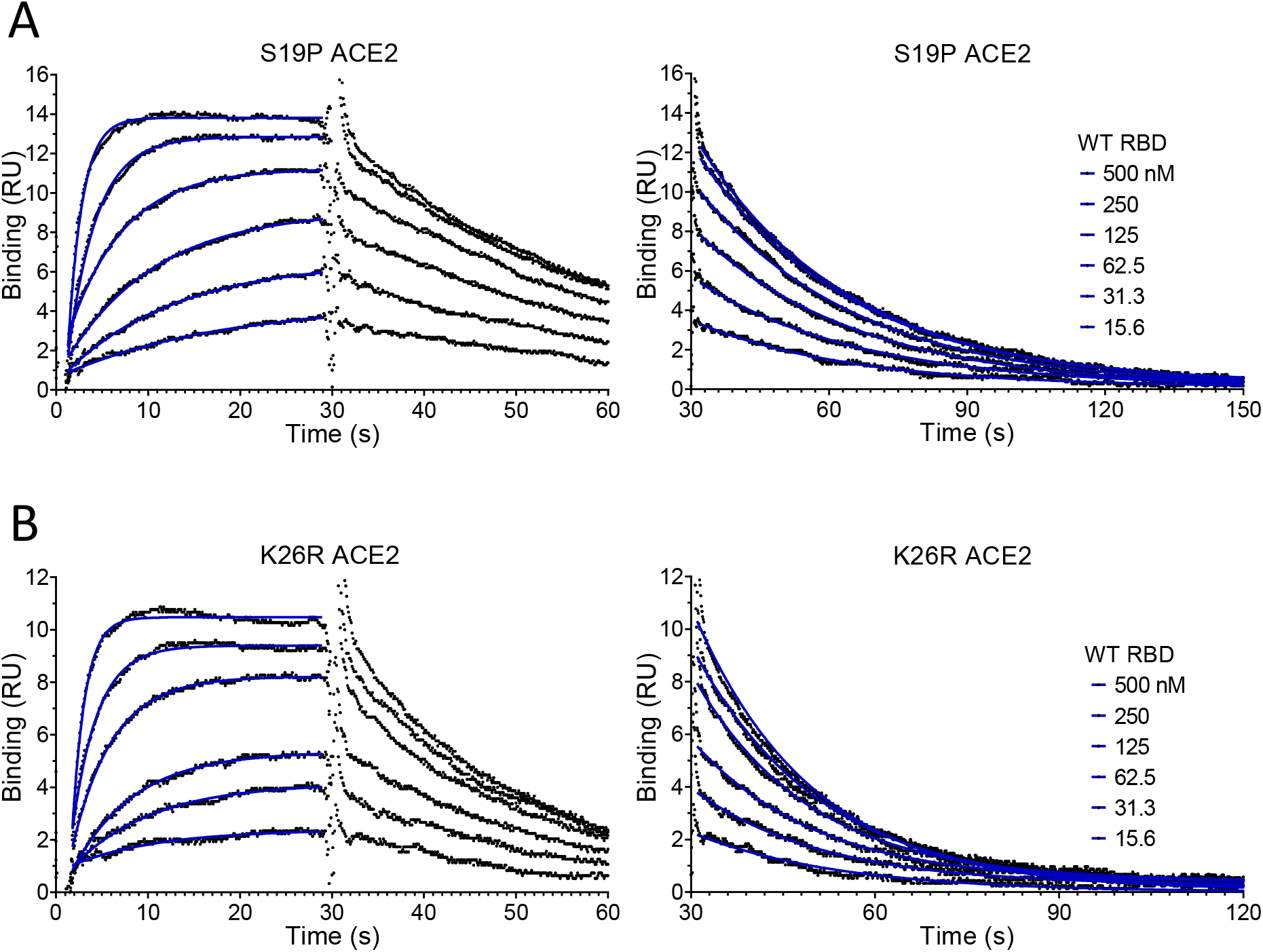
Representative SPR data for WT RBD binding ACE2 variants. Binding traces for the WT RBD injected at different concentrations over the indicated immobilized ACE2 variants. The right panels show an expanded view of the dissociation phase. The blue lines show fits used for determining the k_on_ and k_off_.

## References

Abdiche Y, Malashock D, Pinkerton A, Pons J. 2008. Determining kinetics and affinities of protein interactions using a parallel real-time label-free biosensor, the Octet. Anal Biochem 377:209–217. doi:10.1016/j.ab.2008.03.035

Agius R, Torchala M, Moal IH, Fernández-Recio J, Bates PA. 2013. Characterizing Changes in the Rate of Protein-Protein Dissociation upon Interface Mutation Using Hotspot Energy and Organization. Plos Comput Biol 9:e1003216. doi:10.1371/journal.pcbi.1003216

Cao W, Li T. 2020. COVID-19: towards understanding of pathogenesis. Cell Res 30:367–369. doi:10.1038/s41422-020-0327-4

Cele S, Gazy I, Team C-K, NGS-SA, Jackson L, Hwa S-H, Tegally H, Lustig G, Giandhari J, Pillay S, Wilkinson E, Naidoo Y, Karim F, Ganga Y, Khan K, Bernstein M, Balazs AB, Gosnell BI, Hanekom W, Moosa M-YS, Lessells RJ, Oliveira T de, Sigal A. 2021. Escape of SARS-CoV-2 501Y.V2 from neutralization by convalescent plasma. Nature 1–9. doi:10.1038/s41586-021-03471-w

Darby AC, Hiscox JA. 2021. Covid-19: variants and vaccination. Bmj 372:n771. doi:10.1136/bmj.n771

Davies NG, Edmunds WJ. 2021. Estimated transmissibility and impact of SARS-CoV-2 lineage B.1.1.7 in England. Science. doi:10.1126/science. abg3055

Dejnirattisai W, Zhou D, Supasa P, Liu C, Mentzer AJ, Ginn HM, Zhao Y, Duyvesteyn HME, Tuekprakhon A, Nutalai R, Wang B, Paesen GC, López-Camacho C, Slon-Campos J, Walter TS, Skelly D, Clemens SAC, Naveca FG, Nascimento V, Nascimento F, Costa CF da, Resende PC, Pauvolid-Correa A, Siqueira MM, Dold C, Levin R, Dong T, Pollard AJ, Knight JC, Crook D, Lambe T, Clutterbuck E, Bibi S, Flaxman A, Bittaye M, Belij-Rammerstorfer S, Gilbert S, Carroll MW, Klenerman P, Barnes E, Dunachie SJ, Paterson NG, Williams MA, Hall DR, Hulswit RJG, Bowden TA, Fry EE, Mongkolsapaya J, Ren J, Stuart DI, Screaton GR. 2021. Antibody evasion by the P.1 strain of SARS-CoV-2. Cell. doi:10.1016/j.cell.2021.03.055

Garcia-Beltran WF, Lam EC, Denis KSt, Nitido AD, Garcia ZH, Hauser BM, Feldman J, Pavlovic MN, Gregory DJ, Poznansky MC, Sigal A, Schmidt AG, Iafrate AJ, Naranbhai V, Balazs AB. 2021. Multiple SARS-CoV-2 variants escape neutralization by vaccine-induced humoral immunity. Cell. doi:10.1016/j.cell.2021.03.013

Greaney AJ, Loes AN, Crawford KHD, Starr TN, Malone KD, Chu HY, Bloom JD. 2021a. Comprehensive mapping of mutations in the SARS-CoV-2 receptor-binding domain that affect recognition by polyclonal human plasma antibodies. Cell Host Microbe 29:463–476.e6. doi:10.1016/j.chom.2021.02.003

Greaney AJ, Starr TN, Gilchuk P, Zost SJ, Binshtein E, Loes AN, Hilton SK, Huddleston J, Eguia R, Crawford KHD, Dingens AS, Nargi RS, Sutton RE, Suryadevara N, Rothlauf PW, Liu Z, Whelan SPJ, Carnahan RH, Crowe JE, Bloom JD. 2021b. Complete Mapping of Mutations to the SARS-CoV-2 Spike Receptor-Binding Domain that Escape Antibody Recognition. Cell Host Microbe 29:44–57.e9. doi:10.1016/j.chom.2020.11.007

Hadfield J, Megill C, Bell SM, Huddleston J, Potter B, Callender C, Sagulenko P, Bedford T, Neher RA. 2018. Nextstrain: real-time tracking of pathogen evolution. Bioinformatics 34:4121–4123. doi:10.1093/bioinformatics/bty407

Hasegawa K, Hu C, Nakamura T, Marks JD, Russell SJ, Peng K-W. 2007. Affinity Thresholds for Membrane Fusion Triggering by Viral Glycoproteins▿. J Virol 81:13149–13157. doi:10.1128/jvi.01415-07

Hodcroft EB. 2021. CoVariants: SARS-CoV-2 Mutations and Variants of Interest. https://covariants.org/

Hoffmann M, Arora P, Groß R, Seidel A, Hörnich BF, Hahn AS, Krüger N, Graichen L, Hofmann-Winkler H, Kempf A, Winkler MS, Schulz S, Jäck H-M, Jahrsdörfer B, Schrezenmeier H, Müller M, Kleger A, Münch J, Pöhlmann S. 2021. SARS-CoV-2 variants B.1.351 and P.1 escape from neutralizing antibodies. Cell. doi:10.1016/j.cell.2021.03.036

Hoffmann M, Kleine-Weber H, Schroeder S, Krüger N, Herrler T, Erichsen S, Schiergens TS, Herrler G, Wu N-H, Nitsche A, Müller MA, Drosten C, Pöhlmann S. 2020. SARS-CoV-2 Cell Entry Depends on ACE2 and TMPRSS2 and Is Blocked by a Clinically Proven Protease Inhibitor. Cell. doi:10.1016/j.cell.2020.02.052

Karczewski KJ, Francioli LC, Tiao G, Cummings BB, Alföldi J, Wang Q, Collins RL, Laricchia KM, Ganna A, Birnbaum DP, Gauthier LD, Brand H, Solomonson M, Watts NA, Rhodes D, Singer-Berk M, England EM, Seaby EG, Kosmicki JA, Walters RK, Tashman K, Farjoun Y, Banks E, Poterba T, Wang A, Seed C, Whiffin N, Chong JX, Samocha KE, Pierce-Hoffman E, Zappala Z, O’Donnell-Luria AH, Minikel EV, Weisburd B, Lek M, Ware JS, Vittal C, Armean IM, Bergelson L, Cibulskis K, Connolly KM, Covarrubias M, Donnelly S, Ferriera S, Gabriel S, Gentry J, Gupta N, Jeandet T, Kaplan D, Llanwarne C, Munshi R, Novod S, Petrillo N, Roazen D, Ruano-Rubio V, Saltzman A, Schleicher M, Soto J, Tibbetts K, Tolonen C, Wade G, Talkowski ME, Salinas CAA, Ahmad T, Albert CM, Ardissino D, Atzmon G, Barnard J, Beaugerie L, Benjamin EJ, Boehnke M, Bonnycastle LL, Bottinger EP, Bowden DW, Bown MJ, Chambers JC, Chan JC, Chasman D, Cho J, Chung MK, Cohen B, Correa A, Dabelea D, Daly MJ, Darbar D, Duggirala R, Dupuis J, Ellinor PT, Elosua R, Erdmann J, Esko T, Färkkilä M, Florez J, Franke A, Getz G, Glaser B, Glatt SJ, Goldstein D, Gonzalez C, Groop L, Haiman C, Hanis C, Harms M, Hiltunen M, Holi MM, Hultman CM, Kallela M, Kaprio J, Kathiresan S, Kim B-J, Kim YJ, Kirov G, Kooner J, Koskinen S, Krumholz HM, Kugathasan S, Kwak SH, Laakso M, Lehtimäki T, Loos RJF, Lubitz SA, Ma RCW, MacArthur DG, Marrugat J, Mattila KM, McCarroll S, McCarthy MI, McGovern D, McPherson R, Meigs JB, Melander O, Metspalu A, Neale BM, Nilsson PM, O’Donovan MC, Ongur D, Orozco L, Owen MJ, Palmer CNA, Palotie A, Park KS, Pato C, Pulver AE, Rahman N, Remes AM, Rioux JD, Ripatti S, Roden DM, Saleheen D, Salomaa V, Samani NJ, Scharf J, Schunkert H, Shoemaker MB, Sklar P, Soininen H, Sokol H, Spector T, Sullivan PF, Suvisaari J, Tai ES, Teo YY, Tiinamaija T, Tsuang M, Turner D, Tusie-Luna T, Vartiainen E, Vawter MP, Ware JS, Watkins H, Weersma RK, Wessman M, Wilson JG, Xavier RJ, Neale BM, Daly MJ, MacArthur DG. 2020. The mutational constraint spectrum quantified from variation in 141,456 humans. Nature 581:434–443. doi:10.1038/s41586-020-2308-7

Korber B, Fischer WM, Gnanakaran S, Yoon H, Theiler J, Abfalterer W, Hengartner N, Giorgi EE, Bhattacharya T, Foley B, Hastie KM, Parker MD, Partridge DG, Evans CM, Freeman TM, Silva TI de, Group SC-19 G, Group M of SC-19 G, Angyal A, Brown RL, Carrilero L, Green LR, Groves DC, Johnson KJ, Keeley AJ, Lindsey BB, Parsons PJ, Raza M, Rowland-Jones S, Smith N, Tucker RM, Wang D, Wyles MD, McDanal C, Perez LG, Tang H, Moon-Walker A, Whelan SP, LaBranche CC, Saphire EO, Montefiori DC. 2020. Tracking Changes in SARS-CoV-2 Spike: Evidence that D614G Increases Infectivity of the COVID-19 Virus. Cell 182:812–827.e19. doi:10.1016/j.cell.2020.06.043

Lan J, Ge J, Yu J, Shan S, Zhou H, Fan S, Zhang Q, Shi X, Wang Q, Zhang L, Wang X. 2020. Structure of the SARS-CoV-2 spike receptor-binding domain bound to the ACE2 receptor. Nature 581:215–220. doi:10.1038/s41586-020-2180-5

Lei C, Qian K, Li T, Zhang S, Fu W, Ding M, Hu S. 2020. Neutralization of SARS-CoV-2 spike pseudotyped virus by recombinant ACE2-Ig. Nat Commun 11:2070. doi:10.1038/s41467-020-16048-4

Li W, Zhang C, Sui J, Kuhn JH, Moore MJ, Luo S, Wong S, Huang I, Xu K, Vasilieva N, Murakami A, He Y, Marasco WA, Guan Y, Choe H, Farzan M. 2005. Receptor and viral determinants of SARS‐coronavirus adaptation to human ACE2. Embo J 24:1634–1643. doi:10.1038/sj.emboj.7600640

MacGowan SA, Barton MI, Kutuzov M, Dushek O, van der Merwe PA van der, Barton GJ. 2021. Missense variants in human ACE2 modify binding to SARS-CoV-2 Spike. bioRxiv. doi:10.1101/2021.05.21.445118

Madhi SA, Baillie V, Cutland CL, Voysey M, Koen AL, Fairlie L, Padayachee SD, Dheda K, Barnabas SL, Bhorat QE, Briner C, Kwatra G, Ahmed K, Aley P, Bhikha S, Bhiman JN, Bhorat AE, Plessis J du, Esmail A, Groenewald M, Horne E, Hwa S-H, Jose A, Lambe T, Laubscher M, Malahleha M, Masenya M, Masilela M, McKenzie S, Molapo K, Moultrie A, Oelofse S, Patel F, Pillay S, Rhead S, Rodel H, Rossouw L, Taoushanis C, Tegally H, Thombrayil A, Eck S van, Wibmer CK, Durham NM, Kelly EJ, Villafana TL, Gilbert S, Pollard AJ, Oliveira T de, Moore PL, Sigal A, Izu A, Group N-SGWC. 2021a. Efficacy of the ChAdOx1 nCoV-19 Covid-19 Vaccine against the B.1.351 Variant. New Engl J Med. doi:10.1056/nejmoa2102214

Madhi SA, Baillie V, Cutland CL, Voysey M, Koen AL, Fairlie L, Padayachee SD, Dheda K, Barnabas SL, Bhorat QE, Briner C, Kwatra G, NGS-SA, team W-VC, Ahmed K, Aley P, Bhikha S, Bhiman JN, Bhorat AE, Plessis J du, Esmail A, Groenewald M, Horne E, Hwa S-H, Jose A, Lambe T, Laubscher M, Malahleha M, Masenya M, Masilela M, McKenzie S, Molapo K, Moultrie A, Oelofse S, Patel F, Pillay S, Rhead S, Rodel H, Rossouw L, Taoushanis C, Tegally H, Thombrayil A, Eck S van, Wibmer CK, Durham NM, Kelly EJ, Villafana TL, Gilbert S, Pollard AJ, Oliveira T de, Moore PL, Sigal A, Izu A. 2021b. Safety and efficacy of the ChAdOx1 nCoV-19 (AZD1222) Covid-19 vaccine against the B.1.351 variant in South Africa. medRxiv. doi:10.1101/2021.02.10.21251247

Mahase E. 2021. Covid-19: Where are we on vaccines and variants? Bmj 372:n597. doi:10.1136/bmj.n597

Myszka DG. 1999. Improving biosensor analysis. J Mol Recognit 12:279–284. doi:10.1002/(sici)1099-1352(199909/10)12:5<279::aid-jmr473>3.0.co;2-3

Myszka DG. 1997. Kinetic analysis of macromolecular interactions using surface plasmon resonance biosensors. Curr Opin Biotech 8:50–57. doi:10.1016/s0958-1669(97)80157-7

Pettersen EF, Goddard TD, Huang CC, Couch GS, Greenblatt DM, Meng EC, Ferrin TE. 2004. UCSF Chimera—A visualization system for exploratory research and analysis. J Comput Chem 25:1605–1612. doi:10.1002/jcc.20084

Rambaut A, Holmes EC, O’Toole Á, Hill V, McCrone JT, Ruis C, Plessis L du, Pybus OG. 2020. A dynamic nomenclature proposal for SARS-CoV-2 lineages to assist genomic epidemiology. Nat Microbiol 5:1403–1407. doi:10.1038/s41564-020-0770-5

Rogers TF, Zhao F, Huang D, Beutler N, Burns A, He W, Limbo O, Smith C, Song G, Woehl J, Yang L, Abbott RK, Callaghan S, Garcia E, Hurtado J, Parren M, Peng L, Ramirez S, Ricketts J, Ricciardi MJ, Rawlings SA, Wu NC, Yuan M, Smith DM, Nemazee D, Teijaro JR, Voss JE, Wilson IA, Andrabi R, Briney B, Landais E, Sok D, Jardine JG, Burton DR. 2020. Isolation of potent SARS-CoV-2 neutralizing antibodies and protection from disease in a small animal model. Science 369:956–963. doi:10.1126/science.abc7520

Sabino EC, Buss LF, Carvalho MPS, Prete CA, Crispim MAE, Fraiji NA, Pereira RHM, Parag KV, Peixoto P da S, Kraemer MUG, Oikawa MK, Salomon T, Cucunuba ZM, Castro MC, Santos AA de S, Nascimento VH, Pereira HS, Ferguson NM, Pybus OG, Kucharski A, Busch MP, Dye C, Faria NR. 2021. Resurgence of COVID-19 in Manaus, Brazil, despite high seroprevalence. Lancet 397:452–455. doi:10.1016/s0140-6736(21)00183-5

Sagulenko P, Puller V, Neher RA. 2018. TreeTime: Maximum-likelihood phylodynamic analysis. Virus Evol 4:vex042-. doi:10.1093/ve/vex042

SARS-CoV-2 Variants of concern and variants under investigation - GOV.UK. 2021. https://www.gov.uk/government/publications/covid-19-variants-genomically-confirmed-case-numbers/variants-distribution-of-cases-data

Schreiber G, Fersht AR. 1996. Rapid, electrostatically assisted association of proteins. Nat Struct Biol 3:427–431. doi:10.1038/nsb0596-427

Shang J, Ye G, Shi K, Wan Y, Luo C, Aihara H, Geng Q, Auerbach A, Li F. 2020. Structural basis of receptor recognition by SARS-CoV-2. Nature 581:221–224. doi:10.1038/s41586-020-2179-y

Shu Y, McCauley J. 2017. GISAID: Global initiative on sharing all influenza data –from vision to reality. Eurosurveillance 22:30494. doi:10.2807/1560-7917.es.2017.22.13.30494

Starr TN, Greaney AJ, Hilton SK, Ellis D, Crawford KHD, Dingens AS, Navarro MJ, Bowen JE, Tortorici MA, Walls AC, King NP, Veesler D, Bloom JD. 2020. Deep Mutational Scanning of SARS-CoV-2 Receptor Binding Domain Reveals Constraints on Folding and ACE2 Binding. Cell 182:1295–1310.e20. doi:10.1016/j.cell.2020.08.012

Supasa P, Zhou D, Dejnirattisai W, Liu C, Mentzer AJ, Ginn HM, Zhao Y, Duyvesteyn HME, Nutalai R, Tuekprakhon A, Wang B, Paesen GC, Slon-Campos J, López-Camacho C, Hallis B, Coombes N, Bewley K, Charlton S, Walter TS, Barnes E, Dunachie SJ, Skelly D, Lumley SF, Baker N, Shaik I, Humphries H, Godwin K, Gent N, Sienkiewicz A, Dold C, Levin R, Dong T, Pollard AJ, Knight JC, Klenerman P, Crook D, Lambe T, Clutterbuck E, Bibi S, Flaxman A, Bittaye M, Belij-Rammerstorfer S, Gilbert S, Hall DR, Williams MA, Paterson NG, James W, Carroll MW, Fry EE, Mongkolsapaya J, Ren J, Stuart DI, Screaton GR. 2021. Reduced neutralization of SARS-CoV-2 B.1.1.7 variant by convalescent and vaccine sera. Cell. doi:10.1016/j.cell.2021.02.033

Tegally H, Wilkinson E, Giovanetti M, Iranzadeh A, Fonseca V, Giandhari J, Doolabh D, Pillay S, San EJ, Msomi N, Mlisana K, Gottberg A von, Walaza S, Allam M, Ismail A, Mohale T, Glass AJ, Engelbrecht S, Zyl GV, Preiser W, Petruccione F, Sigal A, Hardie D, Marais G, Hsiao M, Korsman S, Davies M-A, Tyers L, Mudau I, York D, Maslo C, Goedhals D, Abrahams S, Laguda-Akingba O, Alisoltani-Dehkordi A, Godzik A, Wibmer CK, Sewell BT, Lourenço J, Alcantara LCJ, Pond SLK, Weaver S, Martin D, Lessells RJ, Bhiman JN, Williamson C, Oliveira T de. 2021. Emergence of a SARS-CoV-2 variant of concern with mutations in spike glycoprotein. Nature 1–8. doi:10.1038/s41586-021-03402-9

van der Merwe PA van der, Barclay AN. 1996. Analysis of cell-adhesion molecule interactions using surface plasmon resonance. Current opinion in immunology 8:257–261.

V’kovski P, Kratzel A, Steiner S, Stalder H, Thiel V. 2021. Coronavirus biology and replication: implications for SARS-CoV-2. Nat Rev Microbiol 19:155–170. doi:10.1038/s41579-020-00468-6

Volz E, Hill V, McCrone John T., Price A, Jorgensen D, O’Toole Á, Southgate J, Johnson Robert, Jackson B, Nascimento FF, Rey SM, Nicholls SM, Colquhoun RM, Filipe A da S, Shepherd J, Pascall DJ, Shah R, Jesudason N, Li K, Jarrett R, Pacchiarini N, Bull M, Geidelberg L, Siveroni I, Consortium C-U, Koshy C, Wise E, Cortes Nick, Lynch J, Kidd S, Mori M, Fairley DJ, Curran T, McKenna JP, Adams H, Fraser C, Golubchik T, Bonsall D, Moore Catrin, Caddy SL, Khokhar FA, Wantoch M, Reynolds N, Warne B, Maksimovic J, Spellman K, McCluggage K, John M, Beer R, Afifi S, Morgan S, Marchbank A, Price A, Kitchen C, Gulliver H, Merrick I, Southgate J, Guest M, Munn R, Workman T, Connor TR, Fuller W, Bresner C, Snell LB, Charalampous T, Nebbia G, Batra R, Edgeworth J, Robson SC, Beckett A, Loveson KF, Aanensen DM, Underwood AP, Yeats CA, Abudahab K, Taylor BEW, Menegazzo M, Clark G, Smith W, Khakh M, Fleming VM, Lister MM, Howson-Wells HC, Berry Louise, Boswell T, Joseph A, Willingham I, Bird P, Helmer T, Fallon K, Holmes C, Tang J, Raviprakash V, Campbell S, Sheriff N, Loose MW, Holmes N, Moore Christopher, Carlile M, Wright V, Sang F, Debebe J, Coll F, Signell AW, Betancor G, Wilson HD, Feltwell T, Houldcroft CJ, Eldirdiri S, Kenyon A, Davis T, Pybus O, Plessis L du, Zarebski A, Raghwani J, Kraemer M, Francois S, Attwood S, Vasylyeva T, Torok ME, Hamilton WL, Goodfellow IG, Hall G, Jahun AS, Chaudhry Y, Hosmillo M, Pinckert ML, Georgana I, Yakovleva A, Meredith LW, Moses S, Lowe H, Ryan F, Fisher CL, Awan AR, Boyes J, Breuer J, Harris KA, Brown JR, Shah D, Atkinson L, Lee JCD, Alcolea-Medina A, Moore N, Cortes Nicholas, Williams R, Chapman MR, Levett LJ, Heaney J, Smith DL, Bashton M, Young GR, Allan J, Loh J, Randell PA, Cox A, Madona P, Holmes A, Bolt F, Price J, Mookerjee S, Rowan A, Taylor GP, Ragonnet-Cronin M, Nascimento FF, Jorgensen D, Siveroni I, Johnson Rob, Boyd O, Geidelberg L, Volz EM, Brunker K, Smollett KL, Loman NJ, Quick J, McMurray C, Stockton J, Nicholls S, Rowe W, Poplawski R, Martinez-Nunez RT, Mason J, Robinson TI, O’Toole E, Watts J, Breen C, Cowell A, Ludden C, Sluga G, Machin NW, Ahmad SSY, George RP, Halstead F, Sivaprakasam V, Thomson EC, Shepherd JG, Asamaphan P, Niebel MO, Li KK, Shah RN, Jesudason NG, Parr YA, Tong L, Broos A, Mair D, Nichols J, Carmichael SN, Nomikou K, Aranday-Cortes E, Johnson N, Starinskij I, Filipe A da S, Robertson DL, Orton RJ, Hughes J, Vattipally S, Singer JB, Hale AD, Macfarlane-Smith LR, Harper KL, Taha Y, Payne BAI, Burton-Fanning S, Waugh S, Collins J, Eltringham G, Templeton KE, McHugh MP, Dewar R, Wastenge E, Dervisevic S, Stanley R, Prakash R, Stuart C, Elumogo N, Sethi DK, Meader EJ, Coupland LJ, Potter W, Graham C, Barton E, Padgett D, Scott G, Swindells E, Greenaway J, Nelson A, Yew WC, Silva PCR, Andersson M, Shaw R, Peto T, Justice A, Eyre D, Crooke D, Hoosdally S, Sloan TJ, Duckworth N, Walsh S, Chauhan AJ, Glaysher S, Bicknell K, Wyllie S, Butcher E, Elliott S, Lloyd A, Impey R, Levene N, Monaghan L, Bradley DT, Allara E, Pearson C, Muir P, Vipond IB, Hopes R, Pymont HM, Hutchings S, Curran MD, Parmar S, Lackenby A, Mbisa T, Platt S, Miah S, Bibby D, Manso C, Hubb J, Chand M, Dabrera G, Ramsay M, Bradshaw D, Thornton A, Myers R, Schaefer U, Groves N, Gallagher E, Lee D, Williams D, Ellaby N, Harrison I, Hartman H, Manesis N, Patel V, Bishop C, Chalker V, Osman H, Bosworth A, Robinson E, Holden MTG, Shaaban S, Birchley A, Adams A, Davies A, Gaskin A, Plimmer A, Gatica-Wilcox B, McKerr C, Moore Catherine, Williams C, Heyburn D, Lacy ED, Hilvers E, Downing F, Shankar G, Jones H, Asad H, Coombes J, Watkins J, Evans JM, Fina L, Gifford L, Gilbert L, Graham L, Perry M, Morgan M, Bull M, Cronin M, Pacchiarini N, Craine N, Jones R, Howe R, Corden S, Rey S, Kumziene-Summerhayes S, Taylor S, Cottrell S, Jones S, Edwards S, O’Grady J, Page AJ, Wain J, Webber MA, Mather AE, Baker DJ, Rudder S, Yasir M, Thomson NM, Aydin A, Tedim AP, Kay GL, Trotter AJ, Gilroy RAJ, Alikhan N-F, Martins L de O, Le-Viet T, Meadows L, Kolyva A, Diaz M, Bell A, Gutierrez AV, Charles IG, Adriaenssens EM, Kingsley RA, Casey A, Simpson DA, Molnar Z, Thompson T, Acheson E, Masoli JAH, Knight BA, Hattersley A, Ellard S, Auckland C, Mahungu TW, Irish-Tavares D, Haque T, Bourgeois Y, Scarlett GP, Partridge DG, Raza M, Evans C, Johnson K, Liggett S, Baker P, Essex S, Lyons RA, Caller LG, Castellano S, Williams RJ, Kristiansen M, Roy S, Williams CA, Dyal PL, Tutill HJ, Panchbhaya YN, Forrest LM, Niola P, Findlay J, Brooks TT, Gavriil A, Mestek-Boukhibar L, Weeks S, Pandey S, Berry Lisa, Jones K, Richter A, Beggs A, Smith CP, Bucca G, Hesketh AR, Harrison EM, Peacock SJ, Palmer Sophie, Churcher CM, Bellis KL, Girgis ST, Naydenova P, Blane B, Sridhar S, Ruis C, Forrest S, Cormie C, Gill HK, Dias J, Higginson EE, Maes M, Young J, Kermack LM, Hadjirin NF, Aggarwal D, Griffith L, Swingler T, Davidson RK, Rambaut A, Williams T, Balcazar CE, Gallagher MD, O’Toole Á, Rooke S, Jackson B, Colquhoun R, Ashworth J, Hill V, McCrone J.T., Scher E, Yu X, Williamson KA, Stanton TD, Michell SL, Bewshea CM, Temperton B, Michelsen ML, Warwick-Dugdale J, Manley R, Farbos A, Harrison JW, Sambles CM, Studholme DJ, Jeffries AR, Darby AC, Hiscox JA, Paterson S, Iturriza-Gomara M, Jackson KA, Lucaci AO, Vamos EE, Hughes M, Rainbow L, Eccles R, Nelson C, Whitehead M, Turtle L, Haldenby ST, Gregory R, Gemmell M, Kwiatkowski D, Silva TI de, Smith N, Angyal A, Lindsey BB, Groves DC, Green LR, Wang D, Freeman TM, Parker MD, Keeley AJ, Parsons PJ, Tucker RM, Brown R, Wyles M, Constantinidou C, Unnikrishnan M, Ott S, Cheng JKJ, Bridgewater HE, Frost LR, Taylor-Joyce G, Stark R, Baxter L, Alam MT, Brown PE, McClure PC, Chappell JG, Tsoleridis T, Ball J, Gramatopoulos D, Buck D, Todd JA, Green A, Trebes A, MacIntyre-Cockett G, Cesare M de, Langford C, Alderton A, Amato R, Goncalves S, Jackson DK, Johnston I, Sillitoe J, Palmer Steve, Lawniczak M, Berriman M, Danesh J, Livett R, Shirley L, Farr B, Quail M, Thurston S, Park N, Betteridge E, Weldon D, Goodwin S, Nelson R, Beaver C, Letchford L, Jackson DA, Foulser L, McMinn L, Prestwood L, Kay S, Kane L, Dorman MJ, Martincorena I, Puethe C, Keatley J-P, Tonkin-Hill G, Smith C, Jamrozy D, Beale MA, Patel M, Ariani C, Spencer-Chapman M, Drury E, Lo S, Rajatileka S, Scott C, James K, Buddenborg SK, Berger DJ, Patel G, Garcia-Casado MV, Dibling T, McGuigan S, Rogers HA, Hunter AD, Souster E, Neaverson AS, Goodfellow I, Loman NJ, Pybus OG, Robertson DL, Thomson EC, Rambaut A, Connor TR. 2021a. Evaluating the Effects of SARS-CoV-2 Spike Mutation D614G on Transmissibility and Pathogenicity. Cell 184:64–75.e11. doi:10.1016/j.cell.2020.11.020

Volz E, Mishra S, Chand M, Barrett JC, Johnson R, Geidelberg L, Hinsley WR, Laydon DJ, Dabrera G, O’Toole Á, Amato R, Ragonnet-Cronin M, Harrison I, Jackson B, Ariani CV, Boyd O, Loman NJ, McCrone JT, Gonçalves S, Jorgensen D, Myers R, Hill V, Jackson DK, Gaythorpe K, Groves N, Sillitoe J, Kwiatkowski DP, consortium TC-19 GU (COG-Us, Flaxman S, Ratmann O, Bhatt S, Hopkins S, Gandy A, Rambaut A, Ferguson NM. 2021b. Transmission of SARS-CoV-2 Lineage B.1.1.7 in England: Insights from linking epidemiological and genetic data. Medrxiv 2020.12.30.20249034. doi:10.1101/2020.12.30.20249034

Wang Z, Schmidt F, Weisblum Y, Muecksch F, Barnes CO, Finkin S, Schaefer-Babajew D, Cipolla M, Gaebler C, Lieberman JA, Oliveira TY, Yang Z, Abernathy ME, Huey-Tubman KE, Hurley A, Turroja M, West KA, Gordon K, Millard KG, Ramos V, Silva JD, Xu J, Colbert RA, Patel R, Dizon J, Unson-O’Brien C, Shimeliovich I, Gazumyan A, Caskey M, Bjorkman PJ, Casellas R, Hatziioannou T, Bieniasz PD, Nussenzweig MC. 2021. mRNA vaccine-elicited antibodies to SARS-CoV-2 and circulating variants. Nature 1–7. doi:10.1038/s41586-021-03324-6

Washington NL, Gangavarapu K, Zeller M, Bolze A, Cirulli ET, Barrett KMS, Larsen BB, Anderson C, White S, Cassens T, Jacobs S, Levan G, Nguyen J, Ramirez JM, Rivera-Garcia C, Sandoval E, Wang X, Wong D, Spencer E, Robles-Sikisaka R, Kurzban E, Hughes LD, Deng X, Wang C, Servellita V, Valentine H, Hoff PD, Seaver P, Sathe S, Gietzen K, Sickler B, Antico J, Hoon K, Liu J, Harding A, Bakhtar O, Basler T, Austin B, MacCannell D, Isaksson M, Febbo PG, Becker D, Laurent M, McDonald E, Yeo GW, Knight R, Laurent LC, Feo E de, Worobey M, Chiu CY, Suchard MA, Lu JT, Lee W, Andersen KG. 2021. Emergence and rapid transmission of SARS-CoV-2 B.1.1.7 in the United States. Cell. doi:10.1016/j.cell.2021.03.052

Wells JA. 1990. Additivity of mutational effects in proteins. Biochemistry-us 29:8509–8517. doi:10.1021/bi00489a001

WHO Coronavirus (COVID-19) Dashboard. 2021. https://covid19.who.int/

Wölfel R, Corman VM, Guggemos W, Seilmaier M, Zange S, Müller MA, Niemeyer D, Jones TC, Vollmar P, Rothe C, Hoelscher M, Bleicker T, Brünink S, Schneider J, Ehmann R, Zwirglmaier K, Drosten C, Wendtner C. 2020. Virological assessment of hospitalized patients with COVID-2019. Nature 1–10. doi:10.1038/s41586-020-2196-x

Wrapp D, Wang N, Corbett KS, Goldsmith JA, Hsieh C-L, Abiona O, Graham BS, McLellan JS. 2020. Cryo-EM structure of the 2019-nCoV spike in the prefusion conformation. Science 367:1260–1263. doi:10.1126/science.abb2507

Zahradník J, Marciano S, Shemesh M, Zoler E, Chiaravalli J, Meyer B, Rudich Y, Dym O, Elad N, Schreiber G. 2021. SARS-CoV-2 RBD in vitro evolution follows contagious mutation spread, yet generates an able infection inhibitor. Biorxiv 2021.01.06.425392. doi:10.1101/2021.01.06.425392

Zhang J, Cai Y, Xiao T, Lu J, Peng H, Sterling SM, Walsh RM, Rits-Volloch S, Zhu H, Woosley AN, Yang W, Sliz P, Chen B. 2021. Structural impact on SARS-CoV-2 spike protein by D614G substitution. Science eabf2303. doi:10.1126/science.abf2303

Zhang L, Jackson CB, Mou H, Ojha A, Peng H, Quinlan BD, Rangarajan ES, Pan A, Vanderheiden A, Suthar MS, Li W, Izard T, Rader C, Farzan M, Choe H. 2020. SARS-CoV-2 spike-protein D614G mutation increases virion spike density and infectivity. Nat Commun 11:6013. doi:10.1038/s41467-020-19808-4

Zhou D, Dejnirattisai W, Supasa P, Liu C, Mentzer AJ, Ginn HM, Zhao Y, Duyvesteyn HME, Tuekprakhon A, Nutalai R, Wang B, Paesen GC, Lopez-Camacho C, Slon-Campos J, Hallis B, Coombes N, Bewley K, Charlton S, Walter TS, Skelly D, Lumley SF, Dold C, Levin R, Dong T, Pollard AJ, Knight JC, Crook D, Lambe T, Clutterbuck E, Bibi S, Flaxman A, Bittaye M, Belij-Rammerstorfer S, Gilbert S, James W, Carroll MW, Klenerman P, Barnes E, Dunachie SJ, Fry EE, Mongkolspaya J, Ren J, Stuart DI, Screaton GR. 2021. Evidence of escape of SARS-CoV-2 variant B.1.351 from natural and vaccine induced sera. Cell. doi:10.1016/j.cell.2021.02.037

Zhou H, Dcosta BM, Samanovic MI, Mulligan MJ, Landau NR, Tada T. 2021. B.1.526 SARS-CoV-2 variants identified in New York City are neutralized by vaccine-elicited and therapeutic monoclonal antibodies. Biorxiv 2021.03.24.436620. doi:10.1101/2021.03.24.436620

